# Structural basis for Ca^2+^-dependent allosteric modulation of cardiac ryanodine receptor by Ryanozole

**DOI:** 10.64898/2026.05.11.724262

**Authors:** Yuya Otori, Takashi Murayama, Ayuna Tsutsumi, Ryosuke Ishida, Sho Takeuchi, Raymond Burton-Smith, Kazuyoshi Murata, Takashi Sakurai, Hiroyuki Kagechika, Nagomi Kurebayashi, Haruo Ogawa

**Author notes:** Correspondence: (N.K.); (H.O.).

## Abstract

The cardiac ryanodine receptor (RyR2) is a Ca²⁺ release channel essential for excitation–contraction coupling. RyR2 mutations cause severe arrhythmogenic disorders, including catecholaminergic polymorphic ventricular tachycardia (CPVT), through gain-of-function (GOF) effects leading to aberrant Ca²⁺ release. We have recently developed Ryanozole, a potential therapeutic compound for CPVT, which selectively stabilizes RyR2 in a Ca^2+^-dependent manner. Here, we define the mechanism of action of Ryanozole by combining high-resolution cryo-electron microscopy, targeted mutagenesis and functional assays. Ryanozole binds to the interface between the Ca²⁺-binding site and the pore-forming S6 helix, interfering with conformational changes required for Ca²⁺-induced channel opening at low Ca^2+^. We identified key residues for the binding and isoform-specific modulation of Ryanozole. Notably, Ryanozole-bound RyR2 retains its ability to open at high Ca^2+^ via unique conformational changes. These findings provide a structural basis for CPVT-targeted therapy and redefine the paradigm of small molecule-based regulation of large ion channels.

## Introduction

The cardiac ryanodine receptor (RyR2) is a large intracellular Ca²⁺ release channel that plays a central role in myocardial excitation–contraction coupling ^1,2^. During each cardiac cycle, membrane depolarization triggers Ca²⁺ influx through voltage-gated L-type Ca²⁺ channels, which in turn activates RyR2 to release Ca²⁺ from the sarcoplasmic reticulum into the cytosol ^3^. Dysregulation of RyR2 activity has severe physiological consequences. Notably, pathogenic amino acid variants of RyR2 have been linked to catecholaminergic polymorphic ventricular tachycardia (CPVT), a heritable arrhythmogenic disorder characterized by stress-induced ventricular arrhythmias and a high risk of sudden cardiac death ^4–7^. CPVT-associated variants typically enhance the RyR2 channel activity. In response to strong sympathetic stimulation by exercise or emotional stress, these variant RyR2s are further activated, causing spontaneous Ca^2+^ release during diastole. This in turn activates inward sodium-calcium exchanger currents to develop delayed afterdepolarization and triggered activity, leading to arrhythmias ^8,9^.

The major conventional antiarrhythmic drugs currently used for CPVT to include β-blockers, the Na^+^ channel blocker flecainide, and the Ca^2+^ channel blocker verapamil ^10–12^. However, no clinically approved drugs that selectively target RyR2 are currently available for CPVT. An ideal pharmacological strategy for RyR2-targeted drugs must satisfy several stringent criteria imposed by the essential role of RyR2 in cardiac excitation–contraction coupling. These agents should inhibit diastolic Ca^2+^ release with no or minor effects on systolic Ca^2+^ transients for cardiac muscle contraction. They must also exhibit high RyR2 selectivity to avoid off-target inhibition of RyR1 and the associated skeletal muscle dysfunction.

In this context, we have recently established an effective high-throughput screening platform for RyR2 modulators ^13,14^, and developed Ryanozole, a tetrazole compound that selectively stabilizes RyR2 with high affinity in the nanomolar concentration range ^15^. Ryanozole has a unique Ca^2+^-dependent action: it inhibits RyR2 more strongly at lower Ca^2+^ levels ^16^. In CPVT mouse models, Ryanozole suppressed abnormal Ca^2+^ release events at diastole but did not affect action potential-evoked Ca^2+^ transients at systole. Furthermore, it effectively prevented adrenaline-induced arrhythmias and rapidly terminated ongoing spontaneous arrhythmias during daily activity without impairing cardiac conduction or contractility ^16^. These properties distinguish Ryanozole from conventional drugs for CPVT and suggest its potential as a mechanism-based modulator of RyR2 function.

In this study, to gain insights into the molecular basis underlying its mechanism of action, we determined the structures of RyR2 in complex with Ryanozole using cryo-electron microscopy (cryo-EM). We identified the binding site of Ryanozole on the three-dimensional structure of RyR2 at a near-atomic resolution. Based on the structural analysis, we proposed a mode of action for Ca^2+^-dependent modulation and RyR2 specificity. These mechanistic insights were validatedusing extensive targeted mutagenesis and functional assays. Taken together, our findings establish Ryanozole as a structurally and mechanistically defined RyR2-selective modulator and lay the foundation for a rational, mechanism-based approach for correcting pathological Ca²⁺ release in CPVT.

## Results

### Identification of a functional Ryanozole-binding site in RyR2

The RyR2-specific inhibitor Ryanozole features a 3,5-difluorophenyl ring directly linked to a tetrazole ring, which is connected via an N-methylated amide bond to a 4*’*-fluorophenyl ring (**Fig. 1a**) ^15^. The inhibitory effect of Ryanozole on RyR was evaluated using an ER Ca²⁺ assay in HEK293 cells stably expressing both RyR and the Ca²⁺ indicator R-CEPIA1er, in which suppression of RyR channel activity increases ER Ca²⁺ ^13,14,17^ (**Fig. 1b; Extended Data Fig. 1**). Ryanozole selectively suppressed RyR2 (apparent IC50 ≈ 20 nM), with no detectable effects on RyR1 or RyR3 (**Fig. 1c**). A unique property of Ryanozole is that its suppression is greater at lower Ca^2+^ concentrations ^16^. In [³H]ryanodine binding as a functional readout, the apparent IC50 of Ryanozole was 14 nM at low Ca²⁺ concentration (pCa 5.0) but increased to 91 nM at high Ca²⁺ concentration (pCa 3.9) (**Fig. 1d**). Notably, under high Ca²⁺ conditions, Ryanozole did not fully suppress channel opening even at concentrations up to 10 μM (**Fig. 1d**).

**Fig. 1.**
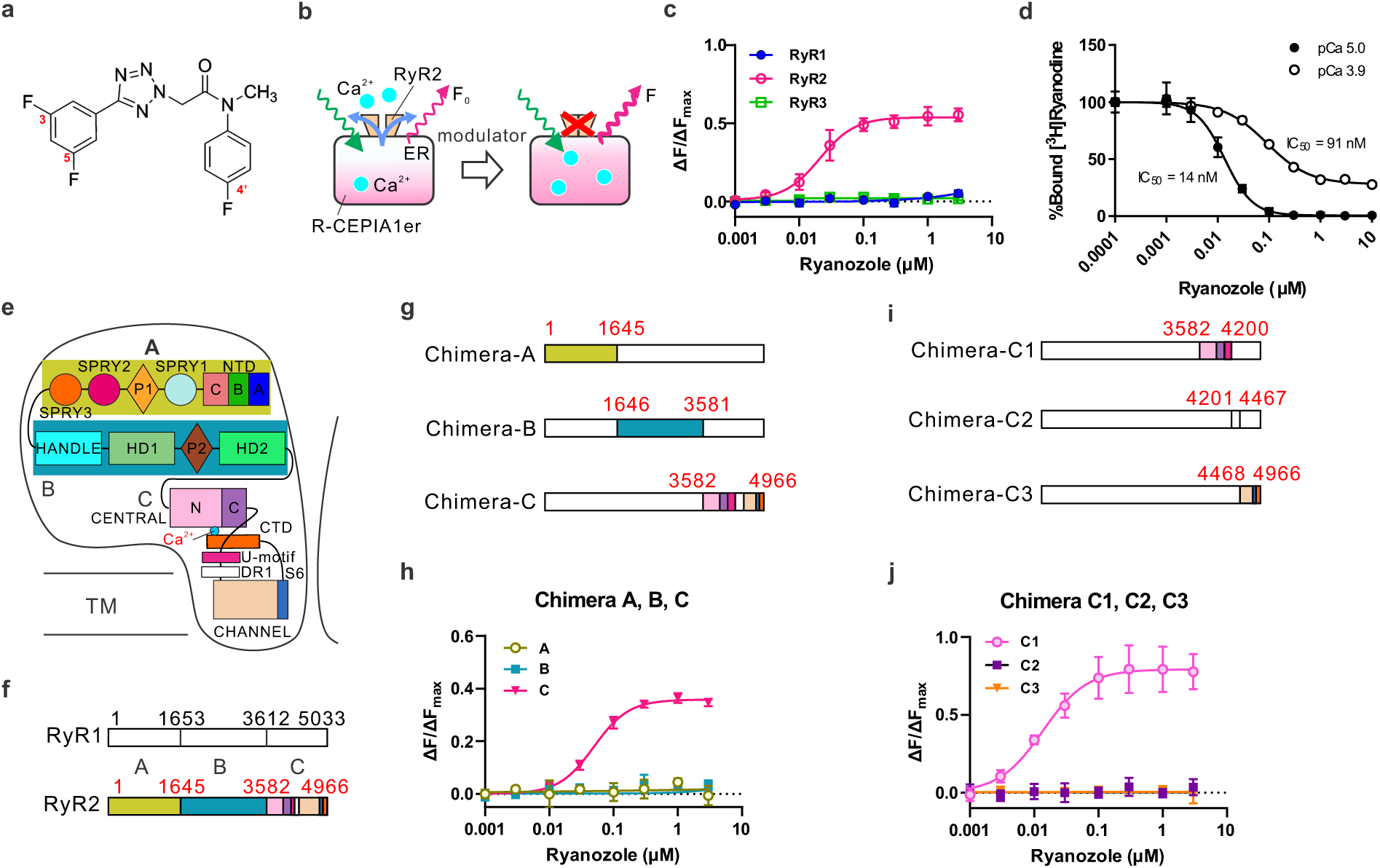
Identification of the Ryanozole binding site. (a) Chemical structure of Ryanozole. (b) Schematic of ER Ca^2+^ measurements used to assess the inhibitory effect of Ryanozole on RyRs. Inhibition of RyR increases ER Ca^2+^ levels, resulting in enhanced R-CEPIA1er fluorescence. (c) Subtype-specific effects of Ryanozole on RyR1, RyR2, and RyR3. Ryanozole selectively inhibits RyR2. (d) Effects of Ryanozole on RyR2 at low (pCa 5.0) and high Ca^2+^ (pCa 3.9) concentrations, assessed by [^3^H]ryanodine binding. Ryanozole inhibits RyR2 more potently under low Ca^2+^ conditions than high Ca^2+^ conditions. (e) Domain organization of RyR. (f) Schematic showing division of RyR1 and RyR2 into three regions based on residue numbering: A (NTD-SPRY3), B (Handle-HD2), and C (Central-CTD). Domain coloring for RyR2 is shown in (e). RyR1-based chimeras generated by replacing each region with the corresponding region from RyR2. (h) Effects of Ryanozole on Chimera-A, -B, and C. Ryanozole inhibits Chimera-C, but not Chimera-A or -B. (i) RyR1-based chimeras in which region C is further subdivided into three domains (C1, C2, and C3). (j) Effects of Ryanozole on Chimera-C1, -C2, and -C3. Ryanozole inhibits Chimera-C1, which contains Central domain and U-motif derived from RyR2.

RyR2 and RyR1 share essentially the same domain organization (**Fig. 1e**) and overall tertiary structure ^18^. To identify the domain responsible for Ryanozole-mediated suppression, we leveraged the differential sensitivities of RyR1 and RyR2 using chimeric constructs. We initially divided full-length RyRs into three regions: A (NTD to SPRY3), B (Handle to HD2), and C (Central domain to CTD) (**Fig. 1f**) and constructed RyR1-based chimeras incorporating the corresponding RyR2 regions (**Fig. 1g**). Among the three chimeras, only Chimera-C exhibited sensitivity to Ryanozole suppression (**Fig. 1h**). We further narrowed down the domain responsible for Ryanozole with three chimeras: Chimera-C1 (Central domain to U-motif), Chimera-C2 (DR1), and Chimera-C3 (Channel domain and beyond) (**Fig. 1i**) and demonstrated that the domain responsible for Ryanozole-mediated suppression exists within Central domain to U-motif (**Fig. 1j**).

### Structural details of the Ryanozole-binding site

To identify the binding site of Ryanozole on RyR2 and the mechanism of its modulation, we performed cryo-EM single-particle analysis. RyR2 was overexpressed in HEK293 cells and purified in complex with SBP-tagged FKBP12.6, as previously described ^19,20^. Cryo-EM datasets were collected with 10 μM Ryanozole under conditions with either 1 mM EGTA or 0.1 mM Ca²⁺, yielding approximately 10,000 movies for each condition. Three-dimensional structures were subsequently reconstructed (**Extended Data Fig. 2**). For higher-resolution structural comparisons, control cryo-EM datasets without Ryanozole were collected under identical conditions (**Extended Data Fig. 3**).

Based on this analysis, we identified three distinct Ryanozole-bound conformations (**Extended Data Fig. 2; Extended Data Fig. 3; Supplementary Table 1; Supplementary Table 2**). To characterize the interaction between Ryanozole and RyR2, we first focused on the closed conformation, in which Ca²⁺ was absent and the channel pore remained closed (Ryanozole/closed; **Fig. 2a**). In this state, Ryanozole binds to the Central domain (**Fig. 2a**), which is consistent with the chimeric analysis (**Fig. 1**). The Central domain is divided into two subdomains, N-Central (residue number: 3605-3987) and C-Central (residue number: 3988-4133), which are separated by Gly3987 ^20^ (**Fig. 2a, right**). N-Central constitutes a Ca^2+^-binding site, and C-Central interacts with U-motif, CTD, and cytoplasmic side of S6 (S6c) to form a tight complex required for channel gating (**Extended Data Fig. 4a**) ^20^. Ryanozole was located within the gap between the N-Central and C-Central domains (**Fig. 2a–b; Extended Data Fig. 4a**). The proximity of this binding site to the Ca²⁺-binding pocket suggests that Ryanozole may modulate the conformational dynamics underlying Ca²⁺-induced activation (**Fig. 2a, right; Extended Data Fig. 4a**). Ryanozole binding was accommodated by pronounced conformational changes in several side chains, including Q3954, Q3974, F4015, and F4016. (**Fig. 2c; Supplementary Movie 1**), without significant main-chain rearrangements in the surrounding N-Central and C-Central domains (**Fig. 2c; Extended Data Fig. 4b**). The 3,5-difluorophenyl ring of Ryanozole is oriented toward the channel pore and extends into the Central domain. At the inner end of this domain, a hydrophobic cluster comprising L3947, M3977, V3978, L3981, and L4002 accommodated the ring, and was likely to be stably positioned (**Fig. 2c**). Together, these findings suggest that the 3,5-difluorophenyl ring enters the binding pocket formed by the N-Central and C-Central domains and is halted by the hydrophobic cluster, with the 4*’*-fluorophenyl ring side serving as the probable entry side (**Fig. 2c**).

**Fig. 2.**
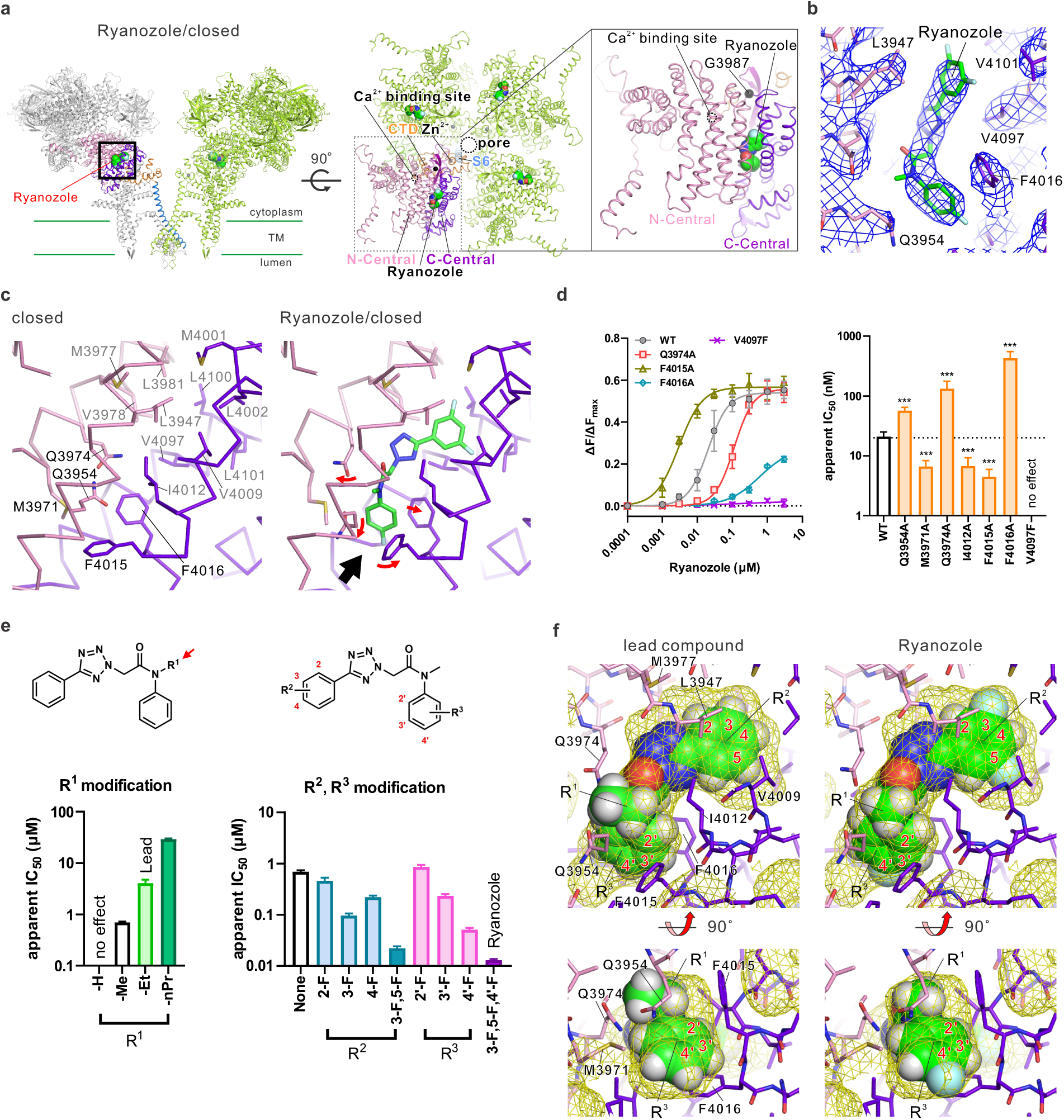
Structure of Ryanozole-bound RyR2. (a) Left, side view of RyR2 in Ryanozole/closed without Ca²⁺. Two opposing subunits of the tetramer are shown, with the Central domain, S6, and CTD of one subunit colored as in Fig. 1e. Bound Ryanozole is depicted as a CPK model. Center, top view of RyR2 near the Ryanozole binding site. Right, enlarged view of the boxed region. (b) Cryo-EM density map of Ryanozole at the binding site, shown in the same orientation as in (a). (c) Structural comparison of residues surrounding the Ryanozole-binding site between the unbound closed state (left) and Ryanozole/closed state (right). Domain coloring follows Fig. 1e. Ryanozole is shown as sticks with carbon atoms in green. Arrows indicate conformational changes associated with Ryanozole binding, and the black arrow indicates a putative entry pathway. (d) Effects of mutations in residues near the Ryanozole binding site on its inhibitory activity. Left, concentration-response curves of Ryanozole for WT, Q3974A, F4015A, F4016A, and V4097F mutants. Right, apparent IC50 values for each mutant. (e) Chemical structure of the lead compound and structure–activity relationship (SAR) analysis of the R¹, R² and R³ substituents. Left, effects of R^1^ substitutions on inhibitory activity. Right, effects of R^2^ and R^3^ substitutions. The R^1^ position was modified by methyl group. Apparent IC50 values were taken from the literature ^15^. (f) Binding pocket of RyR2 visualized after removal of ligand from the Ryanozole/closed structure, overlaid with the lead compound (left) and Ryanozole (right). Ligands are shown as CPK models. The R^1^, R^2^, and R^3^ positions are indicated, and the corresponding ring numbering (Fig. 2e) is shown. Atom colors: carbon (green), nitrogen (blue), oxygen (red), fluorine (cyan), and hydrogen (white).

To assess the contribution of the residues surrounding Ryanozole to ligand binding, alanine substitutions were introduced at seven residues predicted from the Ryanozole-bound structure to participate in ligand interaction: Q3954, M3971, Q3974, I4012, F4015, F4016, and V4097 (**Fig. 2d; Extended Data Fig. 4c**). Q3954A, Q3974A, and F4016A increased the apparent IC50 value, indicating reduced apparent affinity. Because the side chains of Q3954 and Q3974 are located alongside Ryanozole, substitution with alanine is likely to weaken these stabilizing contacts. F4016A markedly diminished the inhibitory effect of Ryanozole, with less than half of the maximally suppressed effect at the highest concentrations tested, highlighting the importance of van der Waals interactions between F4016 and the 4’-fluorophenyl ring of the ligand (**Fig. 2d**). Interestingly, alanine substitutions at the remaining residues (M3971, I4012, and F4015) increased the apparent affinity. F4015 does not directly interact with the ligand and instead appears to partially restrict the optimal positioning of Q3954. Removing this steric constraint by alanine substitution may thus enhance ligand binding. M3971 is located near the entry pathway of the binding pocket. The substitution with alanine may facilitate ligand access to this site. I4012 fits tightly into the concave region formed by the cis-amide configuration of Ryanozole. However, the proximity between the amide oxygen of the ligand and the methyl group of I4012 may be electrostatically unfavorable. Replacement with alanine may alleviate this unfavorable interaction. Furthermore, phenylalanine substitution at V4097, which lies beneath Ryanozole (**Fig. 2c**), completely abolished the effect of Ryanozole (**Fig. 2d**). This suggests that bulky side chain of phenylalanine blocks the Ryanozole-binding pocket and that the binding site of Ryanozole is located exclusively at the N/C-central interface.

Ryanozole was developed by optimizing the systematic structure–activity relationship (SAR) of a lead compound (hit compound in the screening) through modifications at three positions (R^1^, R^2^, and R^3^), which resulted in a more than 300-fold increase in affinity ^15^ (**Fig. 2e**). We interpreted this improvement from a structural perspective by fitting the lead compound into the cavity observed in the Ryanozole-bound structure (**Fig. 2f**). At the R^1^ position, the lead compound contains an ethyl group, whereas Ryanozole has a methyl group. This substitution increased the affinity by approximately fivefold (**Fig. 2e, left**). The ethyl group at R^1^ protrudes beyond the mapped cavity of Ryanozole, causing steric clashes with Q3954 and Q3974 (**Fig. 2f, left**), which may necessitate larger conformational movements of these residues. The R^2^ and R^3^ positions involved modifications of benzene rings, which were unmodified in the lead compound (**Fig. 2e, right**). Introduction of fluorine atoms increased the affinity in a site-specific manner, resulting in an approximately 50-fold enhancement in Ryanozole, which bears 3-F and 5-F substitutions at R^2^ and a 4’-F substitution at R^3^ (**Fig. 2e, right**). Structural fitting showed an additional space around the benzene rings of the lead compound (**Fig. 2f, left**). In contrast, Ryanozole occupied the cavity more completely, exhibiting close steric complementarity with the binding pocket (**Fig. 2f, right**). Taken together, these structural observations provide a mechanistic basis for the SAR trends described above.

### Structural basis of allosteric modulation of RyR2 channel opening by Ryanozole

To address structural basis for modulation of RyR2 channel opening by Ryanozole, we examined two additional conformations: a Ryanozole-bound, Ca²⁺-bound closed conformation (Ryanozole/Ca²⁺/closed; **Fig. 3a**) and a Ryanozole-bound, Ca²⁺-bound open conformation (Ryanozole/Ca²⁺/open; **Fig. 3b**). In both conformations, well-resolved Ryanozole density was observed at the same binding site (**Extended Data Fig. 5a**). The orientations of the side chains surrounding the bound Ryanozole were essentially identical to those observed in the Ryanozole/closed conformation (**Fig. 2c; Extended Data Fig. 5b**), indicating that the Ryanozole-binding mode was preserved across the different channel conformations. Density attributable to Ca²⁺ was observed in both conformations (**Fig. 3a, b**), but appeared relatively stronger in the Ryanozole/Ca²⁺/open conformation than in the Ryanozole/Ca²⁺/closed conformation. Together, these structural observations indicate that Ryanozole can suppress pore opening in the Ca²⁺-bound state, while still allowing channel opening. This interpretation is consistent with the functional data, which demonstrates incomplete suppression of channel activity in the presence of high Ca²⁺ (**Fig. 1d**).

**Fig. 3.**
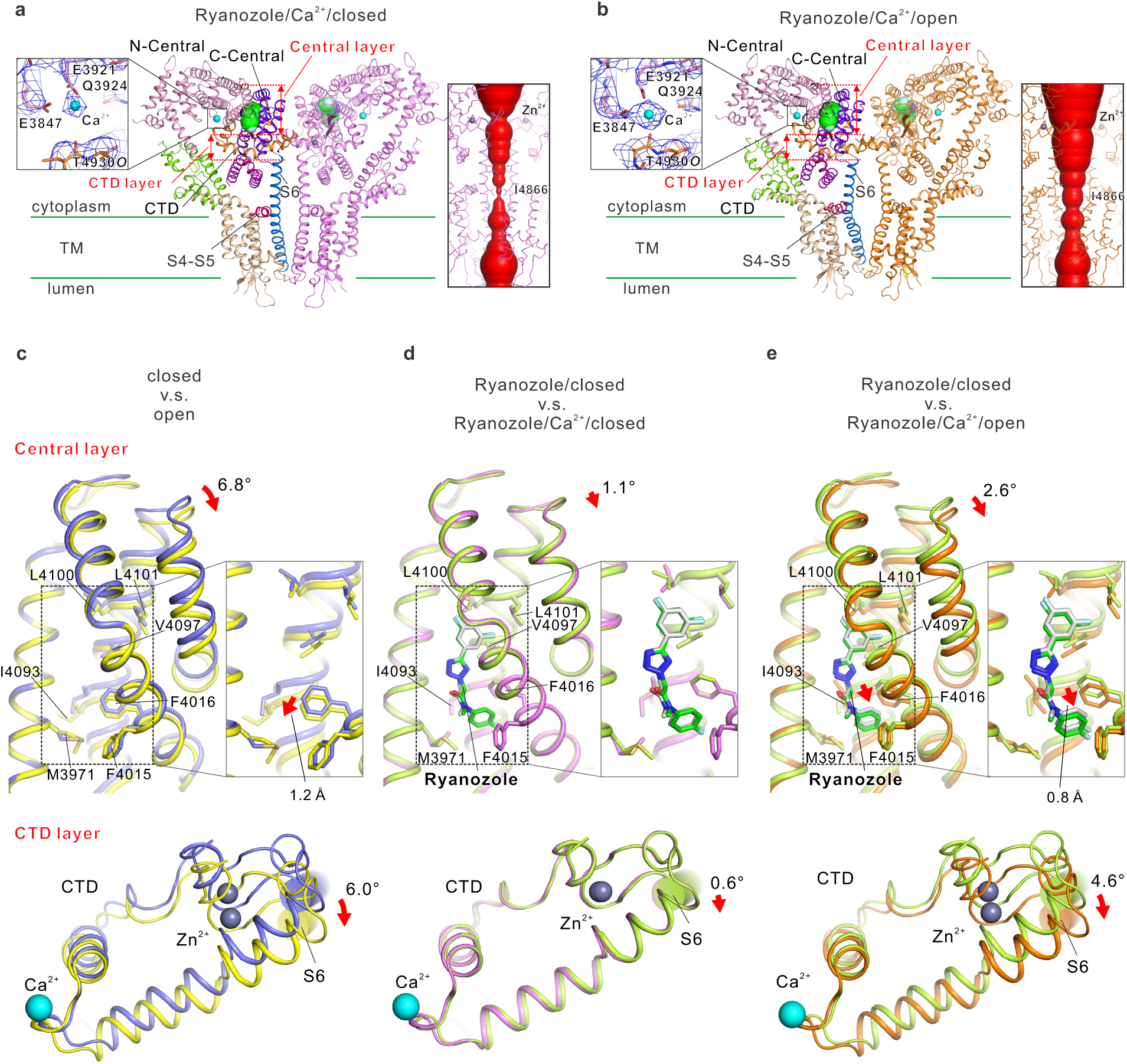
Domain movements in RyR2 upon Ryanozole binding. (a–b) Side views of the RyR2 core domain captured in each alternative conformation with bound Ryanozole. Ryanozole/Ca²⁺/closed conformation (a) and Ryanozole/Ca²⁺/open conformation (b). The left chain is colored as in Fig. 1e. Bound Ryanozole is depicted as a CPK model. Enlarged view of the Ca²⁺ binding site with the Cryo-EM density map shown on the left. Channel pore profiles and pore radii calculated using HOLE ^36^ for each conformation are shown on the right. (c–e) Structural comparisons highlighting Ryanozole-induced domain movements. Top: Relative movement of the C-Central domain with respect to the N-Central domain after N-Central domain fitting; the boxed view on the right shows a magnified view of the dotted boxed area on the left. Bottom: CTD movement after N-Central domain fitting. (c) Closed-to-open transition upon Ca²⁺ binding. Blue, closed; Yellow green, open. (d) Comparison of Ryanozole/closed and Ryanozole/Ca²⁺/closed conformation. Yellow green, Ryanozole/closed conformation; Violet, Ryanozole/Ca²⁺/closed conformation. (e) Comparison of Ryanozole/closed and Ryanozole/Ca²⁺/open conformation. Green, Ryanozole/closed conformation; Orange, Ryanozole/Ca²⁺/open conformation.

We previously showed that Ca²⁺ binding induces separation of the C-Central domain from the N-Central domain, accompanied by a clockwise rotation of t S6c, CTD and the U-motif as a single structural unit, leading to opening of the channel pore ^20^. In the absence of Ryanozole, fitting the C-Central domain in the closed and open states relative to the N-Central domain revealed a clockwise rotation of the C-Central domain by 6.8° (**Fig. 3c, top; Supplementary Movie 2**), accompanied by a corresponding 6.0° rotation of the CTD (**Fig. 3c, bottom; Supplementary Movie 3**). Notably, hydrophobic residues around the Ryanozole-binding pocket on the C-Central domain (F4015, F4016, V4097, L4100, and L4101) (**Fig. 2**) underwent a similar rotational displacement concomitant with rotation of the C-Central domain, and 1.2 Å downward movement of F4016 (**Fig. 3c, top inset**). These residues move collectively as part of the domain rotation.

Next, we applied the same comparative analysis to the three Ryanozole-bound conformations identified in this study. The Ryanozole/closed conformation was nearly identical to the closed conformation (**Extended Data Fig. 5c**), which was consistent with the absence of detectable main-chain rearrangements upon Ryanozole binding (**Fig. 2c; Extended Data Fig. 4b**). Therefore, we compared Ryanozole/Ca^2+^/closed with Ryanozole/closed. In the Ryanozole/Ca^2+^/closed conformation, the Ca²⁺-induced movement of the C-Central domain was severely limited, with rotation confined to the upper portion of the domain (∼1.1°) (**Fig. 3d, top**). This restricted motion was consistent with the steric interference imposed by Ryanozole, together with the associated displacement of F4016 and its neighboring hydrophobic residues (**Fig. 3d, top inset**). Consequently, the Ca²⁺-dependent rotation of the CTD in the region adjacent to S6 was restricted to merely 0.6°, which was insufficient to support the conformational changes required for pore opening (**Fig. 3d, bottom**).

What happened in the open conformation in the presence of Ryanozole? We compared the Ryanozole/Ca^2+^/open conformation with the Ryanozole/Ca^2+^/closed conformation (**Fig. 3e**). The Ca²⁺-dependent rotation of the upper region of the C-Central domain was restricted to approximately 2.6° (**Fig. 3e, top; Supplementary Movie 3**). Instead, the lower region of the C-Central domain accommodated this constrained rotation through a diagonal downward sliding movement of ∼0.8 Å along the bound Ryanozole molecule (**Fig. 3e, top inset**). Consequently, the CTD underwent a 4.6° rotation, which was sufficient to support the S6 rotation and pore opening (**Fig. 3e, bottom; Supplementary Movie 3**). Crucially, the resulting conformation of the C-Central domain in this Ryanozole/Ca²⁺/open state was distinct from that of the open state (**Extended Data Fig. 5c**). Thus, the movement of C-Central domain in the Ryanozole/Ca²⁺/open conformation differed from the canonical closed-to-open transition observed in the absence of Ryanozole (**Fig. 3c, top**).

Together, these structural observations highlight the important contribution of F4016 and the surrounding hydrophobic residues to Ryanozole-mediated modulation of RyR2 gating. This interpretation is supported by the pronounced functional effects of the F4016 mutations observed in our mutagenesis experiments (**Fig. 2d**), suggesting that F4016, together with neighboring hydrophobic residues, forms part of an allosteric network that links Ryanozole binding to channel opening.

### Molecular mechanism underlying RyR2 selectivity of Ryanozole

Finally, we examined the structural basis for the RyR2 selectivity of Ryanozole. In functional study, RyR1-based chimera that contain Central domain derived from RyR2 responded to Ryanozle as RyR2 (**Fig. 1**). This suggests that RyR2 selectivity is simply determined by Central domain. This suggests that RyR2 selectivity is simply determined by Central domain. The amino acid residues in the Ryanozole-binding site within the Central domain were highly conserved among the three isoforms (76.9% similarity for RyR2 vs. RyR1, and 81.2% for RyR2 vs. RyR3), including key residues for Ryanozole binding (L3947, M3977, V3978, L3981, L4002, V4009, I4012, F4015, F4016, V4097, and L4101) (**Extended Data Fig. 6**). Therefore, sequence similarity alone cannot explain the selective inhibition of RyR2 by Ryanozole. We then compared the Ryanozole-binding pocket in available structures for RyR1 ^21^ (PDB ID 8VJJ) and RyR3 ^22^ (PDB ID 9C1E), focusing on the Ryanozole entry gate. Analysis of the cavity surrounding Ryanozole in the bound structure (Ryanozole/closed state) revealed a pocket that was well-suited to accommodate the ligand (**Fig. 4a**). The 3,5-difluorophenyl ring is oriented toward the hydrophobic cluster, whereas the 4 *’*-fluorophenyl group lies adjacent to an external opening positioned between L3967, M3971, and F4086. Therefore, this opening is considered the entry gate for Ryanozole (**Fig. 4a**). The gate, as well as the overall architecture of the pocket accommodating the 3,5-difluorophenyl ring, was preserved even in the Ryanozole-free structure (**Fig. 4b**). However, in the absence of a ligand, the deep portion of the cavity is occluded by the long side chain of Q3974, effectively sealing access to the hydrophobic clusters. Q3974 is a polar residue embedded in a predominantly hydrophobic environment (I4093, I4012, and F4016) and is placed in an energetically strained configuration (**Fig. 4b**). We propose that this local instability primes the ligand-induced rearrangement sites. Upon Ryanozole entry, Q3974 undergoes a substantial conformational shift, vacating the occluded space and permitting the ligand to advance into the deep pocket (**Fig. 2c**). Concomitantly, the side chains along the entry pathway were rearranged (**Fig. 2c; Extended Data Fig. 4b**), collectively stabilizing the bound ligand.

**Fig. 4.**
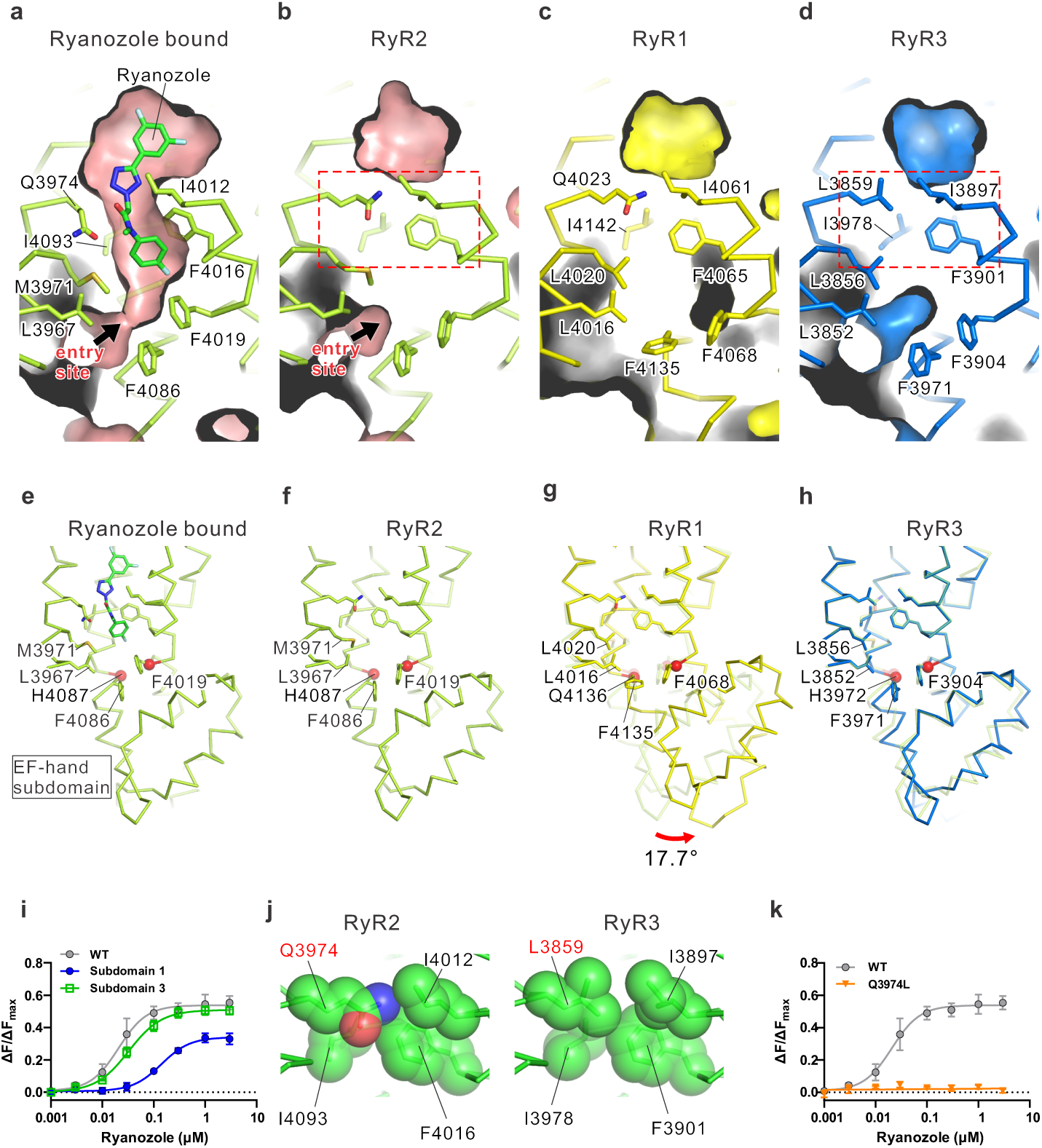
Structural basis for the RyR2 selectivity of Ryanozole. (a-d) Binding pocket of Ryanozole across RyR isoforms. (a) Cavity surrounding Ryanozole in the Ryanozole/closed structure of RyR2. (b-d) Corresponding cavities in the closed state of RyR2 (b), RyR1 (c), and RyR3 (d). (e-h) Ribbon representations of the Central domain around the Ryanozole-binding site, including the EF-hand subdomain in Ryanozole/closed state (e); closed states of RyR2 (f), RyR1 (g), and RyR3 (h). (i) Concentration-dependent effects of Ryanozole on RyR2-based chimeras containing the EF-hand subdomain derived from RyR1 (Subdomain 1) or RyR3 (Subdomain 3). Subdomain 1 largely reduces the affinity for Ryanozole. (j) CPK models of residues within the boxed region shown in (b) and (d). Q3974 in RyR2 is replaced by leucine (L3859) in RyR3. (k) Effect of the Q3974L mutation in RyR2 on concentration-dependent effect of Ryanozole. Q3974L totally abolishes the Ryanozole inhibition.

RyR1 contains a cavity corresponding to the 3,5-difluorophenyl ring–binding site in RyR2; however, no external opening was observed in the region corresponding to the entry gate (formed by L4016, L4020, and F4135) with different directions of the phenylalanine side chain (F4135) (**Fig. 4c**). Upon closer observation of the C-Central structures, we identified distinct conformational differences within the EF-hand subdomain (**Fig. 4e–h**). In RyR1, the EF-hand subdomain is tilted 17.7° clockwise, relative to that in RyR2 or RyR3 (**Fig. 4g**). This reorientation shifts F4135 (corresponding to F4086 in RyR2) toward the gate region, where its side chain projects into the pathway, sterically occluding the passage. Consequently, a functional entry gate cannot be formed in RyR1, which prevents Ryanozole binding. To test this hypothesis directly, we generated chimeric constructs of RyR2 in which the EF-hand subdomain of RyR2 (F4019–H4087) was replaced by the corresponding domain of RyR1 (F4068–Q4136). The chimera showed largely reduced affinity for Ryanozole compared with that of RyR2 (**Fig. 4i**).

RyR3 possesses both a comparable cavity and an external opening in the corresponding entry-gate region (formed by L3852, L3856, and F3971), forming a gate-like structure (**Fig. 4d**). No global changes were observed in the EF-hand subdomain (**Fig. 4h**), with minimal effects of Ryanozole suppression by replacing the EF-hand subdomain with the corresponding RyR3 domain (F3904–H3971) (**Fig. 4i**). In RyR3, the residue corresponding to Q3974 in RyR2 was changed to L3859 (**Fig. 4d**). Unlike the polar glutamine, leucine forms stable interactions with neighboring hydrophobic residues (I3897, F3901, and I3978), thereby stabilizing its side-chain conformation within the pocket (**Fig. 4j**). This hydrophobic network likely restricts the conformational flexibility required for cavity opening (**Fig. 4d**). Consistently, substitution of Q3974 with leucine in RyR2 nearly abolished the suppressive activity of Ryanozole (**Fig. 4k**). Together, these results support an isoform-specific, gate-controlled, induced-fit mechanism in which ligand access and deep-pocket engagement depend on the presence of a conformationally permissive occluding residue.

## Discussion

In this study, we defined a discrete and mechanistically informative binding site for Ryanozole on RyR2, providing insights into how Ca²⁺ sensing is coupled to channel gating, and how this process can be pharmacologically modulated. Structural analysis revealed that Ryanozole binds uniquely at the boundary between the N-Central and C-Central subdomains of RyR2 within the tetrameric channel (**Fig. 2**; **Fig. 3**). This region lies directly above the C-terminal domain (CTD), an elongated structural element that plays a central role in Ca²⁺ coordination (**Fig. 3**). On the N-Central side, the CTD interacts closely with the Central domain to form a composite Ca²⁺-binding environment, whereas on the C-Central side the CTD is mechanically linked to the pore-lining S6 helix. This architecture positions the Ryanozole-binding site precisely along the structural axis that transmits Ca²⁺-binding information from the cytosolic domains to the transmembrane gate (**Fig. 3**).

In RyR2 channel opening, Ca²⁺ binds between the N-Central domain and the CTD, inducing a clockwise rotation of the CTD. This rotation drives a corresponding clockwise rotation of the S6 helix, which forms the channel pore, thereby opening RyR2. The CTD rotation also separates the N-Central and the C-terminal domain, accompanied by an independent clockwise rotation of the C-Central domain relative to the N-Central domain (**Fig. 5a, top**). Ryanozole binds to the cleft between the N-Central and C-Central domains (**Fig. 5a, lower left**). Stabilized by surrounding hydrophobic residues near F4016, Ryanozole restricts the clockwise rotation of the C-Central domain even in the presence of Ca²⁺, thereby preventing CTD-coupled rotation of S6 and channel opening (**Fig. 5a, lower middle**). This may effectively stabilize the channel in the closed state especially at low Ca^2+^ condition, in which population of open channel is low. We also identified the open state structure in the presence of Ryanozole, but via an alternative mechanism distinct from the canonical open state (**Fig. 5a, top right**). In this mode, the upper portion of the C-Central domain undergoes a slight clockwise rotation upon Ca²⁺ binding, while the lower portion glides downward along the Ryanozole-binding surface. The combination of these movements transmits rotation to S6 through the CTD, allowing channel opening despite Ryanozole binding (**Fig. 5a, lower right**). This state may be converted from the Ca^2+^/Ryanozole/closed state to some extent and is likely to occur at high Ca^2+^ condition, where Ca^2+^-binding site is highly occupied. Thus, Ryanozole represents a mechanistically ideal compound that, at low Ca²⁺ concentrations, halts the rotation of S6 just prior to the Ca²⁺-induced conformational cascade ^20^, while at high Ca²⁺ concentrations, it permits a pseudo-rotation of S6 (**Fig. 5a, lower right**). This enables its unique Ca^2+^-dependent modulation. In addition, prevention of Ca²⁺-induced channel activation by Ryanozole reasonably explains that it essentially stabilizes all the gain-of-function mutants ^16^.

**Fig. 5.**
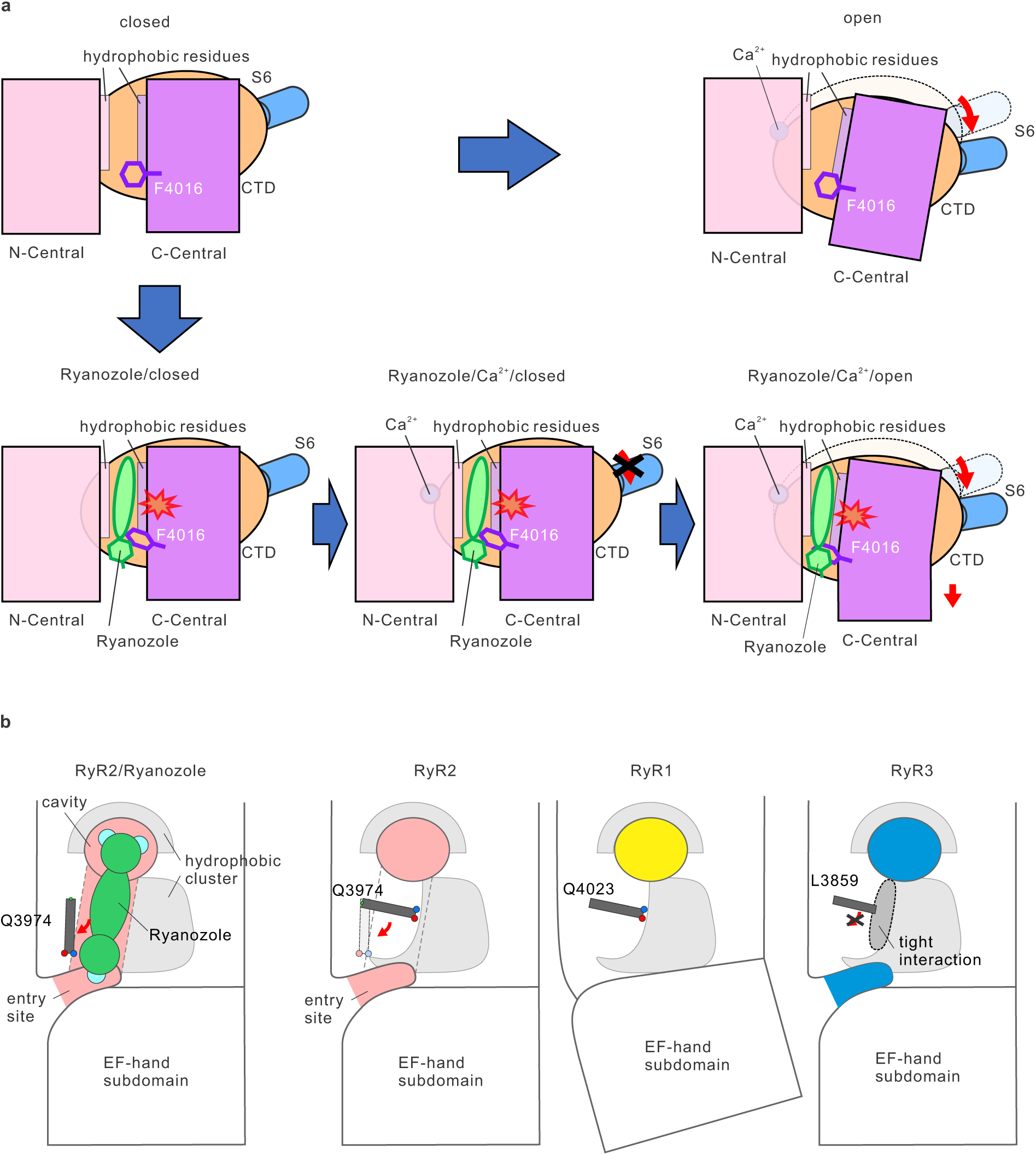
Schematic model of the Ryanozole-mediated modulation of RyR2 gating. (a) Conformational transitions of RyR2 in the presence and absence of Ryanozole. Schematic representations show a single chain comprising the N-Central (pink) and C-Central (purple) domains, the CTD (brown), and the S6 helix (blue). In the absence of Ryanozole (top row), binding of Ca^2+^ induces a large rotation of the C-Central domain, triggering CTD movement and S6 pore opening (from top left to top right). In the presence of Ryanozole (bottom row), it binds at the interface between the N-Central and C-Central domains, stabilized by a hydrophobic cluster including F4016 (purple) (bottom left). Upon Ca^2+^ binding, the steric presence of Ryanozole restricts the canonical rotation of the C-Central domain to stabilize the channel at the closed state (bottom middle). To accommodate this constraint, the C-Central domain undergoes a diagonal downward sliding movement along the Ryanozole molecule (bottom right, red arrow), which allows for the rotation of the CTD and subsequent S6 opening. (b) Structural basis for the isoform-specific selectivity of Ryanozole. Detailed views of the Ryanozole-binding pocket across RyR isoforms illustrate the mechanisms of binding and exclusion. In the Ryanozole-bound RyR2 (far left), Ryanozole is stably accommodated within the binding cavity. In RyR2 without Ryanozole (middle left), although the cavity and entry site are present, the internal passage is obstructed by the side chain of Q3974. In RyR1 (middle right), the entry site is absent due to the distinct orientation of the EF-hand subdomain compared to RyR2, and the internal passage remains blocked by Q4023, collectively preventing Ryanozole binding. In RyR3 (far right), while the potential entry site and the cavity for the 3,5-difluorophenyl ring exist, the passage is occluded by L3859; which forms a tight interaction with neighboring hydrophobic residues, creating a steric barrier that precludes Ryanozole access.

The isoform specificity of Ryanozole provides critical insights into the structural determinants governing the access to its binding pathways. In RyR2, the entry gate is positioned between the EF-hand subdomain at the tip of the Central domain and the Central domain itself, forming a gated access pathway for Ryanozole (**Fig. 5b, left**). From this gate, a continuous cavity extends deep into the Central domain, along which the ligand progresses until the 3,5-difluorophenyl ring is embedded within a hydrophobic cluster located at the distal end of the pocket. However, in the ligand-free state, this cavity is sealed by the long side chain of Q3974 (**Fig. 5b, second from the left**), thereby preventing access to the deep binding site. Notably, Q3974 is a polar residue surrounded by hydrophobic residues, placing it in an energetically unfavorable environment (**Fig. 5b, second from the left**). We propose that this strained configuration is destabilized upon Ryanozole entry, causing Q3974 to undergo a substantial conformational shift that vacates the occluded space and opens the cavity (**Fig.5b, left**). This ligand-triggered rearrangement enables accommodation of Ryanozole and stabilizes the final bound state, which is consistent with an induced-fit mechanism that couples gate opening with deep-pocket engagement. In contrast, in RyR1, the EF-hand subdomain adopts a markedly different orientation relative to the Central domain (**Fig. 5b, third from the left**). Consequently, the entry gate formed between the EF-hand and Central domains in RyR2 is absent, precluding Ryanozole binding (**Fig. 5b, third from the left**). In RyR3, the EF-hand subdomain is oriented similarly to that in RyR2, and an entry gate-like structure is formed (**Fig. 5b, right**). However, the residue corresponding to Q3974 in RyR2 (L3859 in RyR3) forms stable hydrophobic interactions with the surrounding residues. This stabilizing interaction restricts the side-chain mobility and prevents the conformational rearrangement required for cavity opening, thereby obstructing ligand entry and explaining the lack of Ryanozole binding (**Fig. 5b, right**).

Consistent with this structural framework, SAR analyses further supported the mechanism of Ryanozole binding ^15^. Optimization from the lead compound enhanced its affinity through substitutions that improved steric complementarity within the binding pocket, particularly involving the 3,5-difluorophenyl moiety. These modifications aligned with the structural observation that the ligand was tightly accommodated within a hydrophobic cluster at the distal end of the cavity, thereby reinforcing our model. These structural and SAR insights will be a powerful guide for the rational optimization of Ryanozole derivatives toward improved pharmacokinetic and drug-like properties.

Recent studies have identified several small-molecule compounds that stabilize RyR2, including flecainide, carvedilol, EL20, Rycals, and unnatural verticilide enantiomers ^23,24^. Among these, Rycals, such as ARM210, have been reported to bind to the P1 domain and stabilize interdomain interactions ^25,26^. However, their limited isoform selectively and distinct binding sites suggests mechanisms that differ fundamentally from that of Ryanozole ^27–29^. Further structural characterization of RyR2-targeting compounds will be essential for understanding the diverse mechanisms by which small molecules modulate channel gating and for guiding the development of more selective therapeutics.

Finally, we revealed the structure of RyR2 bound to Ryanozole and elucidated its modulatory mechanism, which was further supported by functional analyses that validated the structural model. Together with its previously demonstrated efficacy in disease model mice, these results suggest that Ryanozole is a promising candidate therapeutic agent for CPVT.

## Methods

### Construction of mutant RyR2 cDNA expression vectors and generation of stable HEK293 cell lines

RyR1/RyR2 and RyR2/RyR3 chimeric constructs were generated using the In-Fusion cloning method with the overlapping primers listed in Supplementary Table 3. Point mutations in RyR2 were introduced by inverse PCR using the primers listed in Supplementary Table 4. All mutations were verified using DNA sequencing. The resulting cDNA was subcloned into a pcDNA5/FRT/TO vector. Stable HEK293 cell lines expressing the mutant RyR2 were established using the Flp-In T-REx system (Thermo Fisher Scientific) according to the manufacturer’s instructions. Clones exhibiting appropriate protein expression levels were selected by western blotting and used for subsequent experiments.

### Expression of RyR2 and preparation of microsomes

The expression of mouse wild-type (WT) RyR2 was performed following the established protocols ^20^. Briefly, HEK293 cells stably expressing mouse WT RyR2 were grown in eighty 150-mm dishes. Upon reaching 70–80% confluency, protein expression was induced with 2 µg/mL doxycycline (Sigma) for 48 h. Cells were then harvested, washed with ice-cold phosphate-buffered saline (PBS) (Gibco), and microsomes were prepared as described by Inesi et al ^30^. The cell pellet was resuspended in 60 mL of 10 mM NaHCO₃ containing a protease inhibitor cocktail (Fujifilm Wako) and subjected to nitrogen cavitation for 30 min at 1,000 psi. The resulting suspension was diluted with 60 mL of 0.6 M sucrose, 0.3 M KCl, 40 mM MOPS-Na (pH 7.4), and protease inhibitor cocktail, followed by centrifugation at 1,000 × *g* for 10 min. The supernatant was supplemented with 30 mL of 2.4 M KCl, 0.3 M sucrose, 20 mM MOPS-Na (pH 7.4), and protease inhibitors, and centrifuged at 10,000 × *g* for 20 min. The resulting supernatant was then ultracentrifuged at 100,000 × *g* for 30 min. The microsomal pellet was resuspended in 60 mL of 0.6 M KCl, 0.3 M sucrose, 20 mM MOPS-Na (pH 7.4), and protease inhibitors and ultracentrifuged again. Finally, the pellet was resuspended in 12 mL of 0.3 M sucrose, 20 mM MOPS-Na (pH 7.4), and protease inhibitors, rapidly frozen in liquid nitrogen, and stored at −80°C until further use.

### Purification of RyR2

Purification was performed using affinity chromatography with SBP-tagged FKBP12.6 ^19,20^. Microsomes were solubilized in a solution with 2% (w/v) CHAPS (Dojindo) and 1% (w/v) soybean lecithin (Avanti Polar Lipids) in 1 M NaCl, 20 mM MOPS (pH 7.4), 2 mM dithiothreitol, and a protease inhibitor cocktail for 30 min on ice. After centrifugation at 100,000 × *g* for 30 min at 4°C, the resulting supernatant was diluted four-fold with 20 mM MOPS (pH 7.4), 2 mM dithiothreitol, and protease inhibitors, passed through a 0.45-μm filter, and loaded onto a pre-equilibrated 5-mL StrepTrap HP column (GE Healthcare, Chicago, IL) containing the immobilized SBP-FKBP12.6 fusion protein. The column was washed with 10 column volumes of (1) wash buffer (20 mM MOPS pH 7.4, 2 mM dithiothreitol, and 0.3 M sucrose) containing 0.2 M NaCl and 0.25% (w/v) CHAPS, followed by (2) wash buffer containing 0.5 M NaCl and 0.015% (w/v) Tween-20 (Sigma-Aldrich). The SBP-FKBP12.6–RyR2 complex was eluted with wash buffer supplemented with 2.5 mM D-desthiobiotin (Iba Life Sciences). After verifying purity using SDS–PAGE, the eluate was rapidly frozen in liquid nitrogen and stored at −80°C until further use.

### Three-dimensional structural analysis by cryo-EM

The purified RyR2 sample was buffer-exchanged and concentrated to ∼5 mg/mL using an Amicon Ultra 100k device (Millipore, Burlington, MA) in buffer containing 0.5 M NaCl, 20 mM MOPS-Na (pH 7.4), 2 mM dithiothreitol, and 0.015% (w/v) Tween-20, supplemented with either 1 mM EGTA or 100 µM CaCl₂. For Ryanozole-bound samples, Ryanozole was added at a final concentration of 10 μM and incubated at 4°C for 30 min. Control samples without Ryanozole were prepared using the same EGTA or Ca²⁺ conditions. The samples were applied to Quantifoil Au grids (R1.2/1.3, 300 mesh) (Quantifoil), blotted for 4 s at 100% humidity and 6°C using a Vitrobot Mark IV (Thermo Fisher), and plunge-frozen in liquid ethane.

The grids were initially screened for ice thickness and particle quality using a Glacios cryo-EM (Thermo Fisher Scientific, Waltham, MA) operated at 200 kV and equipped with a Falcon 4 detector (Kyoto University, Japan). The selected grids were subsequently imaged using a Krios G4 cryo-EM (Thermo Fisher Scientific, Waltham, MA, USA) operating at 300 kV and equipped with a Selectris X energy filter (slit width 10 eV) and a Falcon 4i direct electron detector (Thermo Fisher Scientific) in the electron counting mode. Data were collected at a nominal magnification of 130,000×, corresponding to a calibrated pixel size of 0.92 Å/pixel (NIPS, Japan), with a defocus range of −0.5 to −2.0 µm.

Image processing and 3D reconstruction were performed using CryoSPARC 4.6 ^31^. Density-map visualization was performed using UCSF ChimeraX ver. 1.8 ^32^. Model building was conducted in COOT ver. 0.9.8.5 ^33^, and structural refinement was performed using Phenix ver. 1.19.2 ^34,35^. All figures were prepared using PyMOL (The PyMOL Molecular Graphics System; http://www.pymol.org). Pore radii along the ion conducting pathway were calculated using HOLE^36^.

### Synthesis of Ryanozole and related compounds

Ryanozole and its analogs were synthesized and prepared according to previously described procedures. The name of Ryanozole in the literature is TMDJ-035, and the names of its analogs are TMDJ-001–003, TMDJ-015–018, and TMDJ-022–024. ^15^.

### Evaluation of the inhibitory effects of Ryanozole on WT and mutant RyRs in HEK293 cells

Time-lapse measurements of ER Ca^2+^ were performed using a FlexStation 3 fluorometer (Molecular Devices) as previously described ^14,17^. This assay is based on the restoration of reduced ER Ca^2+^ levels caused by spontaneous Ca^2+^ release through the expressed RyRs. Inhibition of RyR-mediated Ca^2+^ release increases ER Ca^2+^ content, which can be monitored as an increase in the fluorescence of the ER-targeted indicator R-CEPIA1er. HEK293 cells were seeded into 96-well flat, clear-bottom black microplates (655090, Greiner) in culture medium supplemented with R-CEPIA1er-encoding baculovirus on day 1. RyR expression was induced by adding doxycycline (2 μg/mL) on day 3. ER Ca^2+^ levels were measured 24 h after induction (day 4). For some mutant RyR2s exhibiting relatively low basal ER Ca^2+^ depletion, doxycycline was added on day 2 instead of day 3 to ensure sufficient reduction in basal ER Ca^2+^ before measurement on day 4. This adjustment was introduced to standardize assay sensitivity across RyR variants, as the detection of inhibitor effects requires partially depleted ER Ca^2+^ stores. Similarly, RyR1, RyR3, and RyR1-RyR2 chimera A exhibited minimal ER Ca^2+^ depletion, regardless of the induction duration. For these, measurements were performed in HEPES–Krebs solution supplemented with 2–4 mM caffeine to establish measurable ER Ca^2+^ depletion ^14,17^.

Before measurement, the culture medium was replaced with 81 μL HEPES–Krebs solution containing 140 mM NaCl, 5 mM KCl, 2 mM CaCl2, 1 mM MgCl2, 11 mM glucose, and 10 mM HEPES (pH 7.4). R-CEPIA1er fluorescence was excited at 560 nm and emission at 610 nm was recorded every 10 s for 260 s. At 60 s, 54 μL of the compound solution (0.0025–7.5 μM in HEPES–Krebs solution) was added to achieve final concentrations of 0.001–3 μM. The fluorescence intensities at the initial 60 s (F0) and final 60 s (F) were quantified. The maximal fluorescence (Fmax) of R-CEPIA1er was determined by applying an Fmax cocktail containing 20 mM Ca^2+^, 20 μM ionomycin and 10 mM caffeine. Here, caffeine was included to facilitate Ca^2+^ permeation through RyRs; under conditions where the external Ca²⁺ concentration exceeds that in the ER lumen, RyR activation promotes Ca²⁺ influx into the ER, resulting in net Ca²⁺ accumulation in the ER ^37^. The effects of Ryanozole in each mutant cell line were expressed as Δ F/ Δ Fmax, calculated as (F – F0)/(Fmax – F0) (**Extended Data Fig. 1**). Normalization to Fmax minimized the variability arising from the differences in basal ER Ca^2+^ content among RyR isoforms and mutants. The concentration-dependent effects of Ryanozole and its precursor compounds on WT and mutant RyRs were analyzed by fitting the dose-response curves using Prism version 11 (GraphPad Software, Inc., La Jolla, CA, USA). from which apparent IC50 values were obtained.

### [^3^H]Ryanodine binding assay

[^3^H]Ryanodine binding assays were performed as described previously ^16,20^. Briefly, microsomes prepared from HEK293 cells expressing wild-type RyR2 were incubated for 1 h at 25 °C with 5 nM [³H]ryanodine (PerkinElmer) in a reaction buffer containing 0.17 M NaCl, 20 mM MOPSO-Na (pH 7.0), 2 mM dithiothreitol, 1 mM AMP, and 1 mM MgCl2, in the presence of 0–10 µM Ryanozole. Free Ca^2+^ concentrations were adjusted to pCa 5.0 or 3.9 using 10 mM EGTA and calculated with WEBMAXC STANDARD (https://somapp.ucdmc.ucdavis.edu/pharmacology/bers/maxchelator/webmaxc/webmaxcS.htm)^38^. [^3^H]ryanodine binding at each Ryanozole concentration was normalized to that obtained in the absence of Ryanozole.

## Data availability

Atomic coordinates and cryo-EM density maps for the global structures and core domain (obtained via local refinement) have been deposited in the PDB and EMDB with the following accession codes: 26IP and EMD-80684 (overall closed state), 26IU and EMD-80689 (core domain closed state); 26IO and EMD-80683 (overall open state), 26IT and EMD-80688 (core domain open state); 26IS and EMD-80687 (overall Ryanozole/closed conformation), 26IX and EMD-80692 (core domain Ryanozole/closed conformation); 26IR and EMD-80686 (overall Ryanozole/Ca^2+^/closed conformation), 26IW and EMD-80691 (core domain Ryanozole/Ca^2+^/closed conformation); 26IQ and EMD-80685 (overall Ryanozole/Ca^2+^/open conformation), 26IV and EMD-80690 (core domain Ryanozole/Ca^2+^/open conformation).

## Acknowledgements

We thank Dr. Sugita and Dr. Kimura at the Institute for Life and Medical Sciences, Kyoto University, for their assistance with sample screening using the Glacios cryo-EM and Ikue Hiraga and staff at the Laboratory of Proteomics and Biomolecular Science, Research Support Center, Juntendo University Graduate School of Medicine, for technical assistance. This study was partly supported by the Japan Society for the Promotion of Sciences KAKENHI (grant numbers 21H02411 and 24K02164 to H.O., 22K15244 to R.I., 25K02443 to T.M., and 22K06652 to N.K.), the Platform Project for Supporting Drug Discovery and Life Science Research (Basis for Supporting Innovative Drug Discovery and Life Science Research [BINDS] grant number JP21am0101080 to H.O. and T.M., JP25ama121005 to K.M., and JP25ama121043 to R.I. and H.K.), oint research program of National Institute for Physiological Sciences (grant number 25NIPS209 to H.O.), JST SPRING (grant number JPMJSP2110 to Y.O. and A.T.), and the Vehicle Racing Commemorative Foundation (6516 to N.K.). A part of this research was based on the Cooperative Research Project of the Research Center for Biomedical Engineering.

## Author contributions

T.M., N.K., and H.O. conceived and designed the study. Y.O. and A.T. performed cell culture experiments. Y.O. purified the proteins. Y.O. and H.O. prepared the cryo-EM grids. Y.O., R.B-S., and K.M. acquired the EM images. Y. O., S.T., and H.O. processed the images. Y.O. and H.O. performed model building and refinement. T.M. and N.K. generated chimeric and mutant constructs and performed the functional analysis. R.I. and H.K. synthesized Ryanozole and its related compounds. Y.O., T.M., N.K., T.S., and H.O. interpreted the data. N.K. and H.O supervised the study. Y.O., N.K., T.M., and H.O. wrote the manuscript with input from all the authors.

## Extended Data Fig. legends

**Extended Data Fig. 1.**
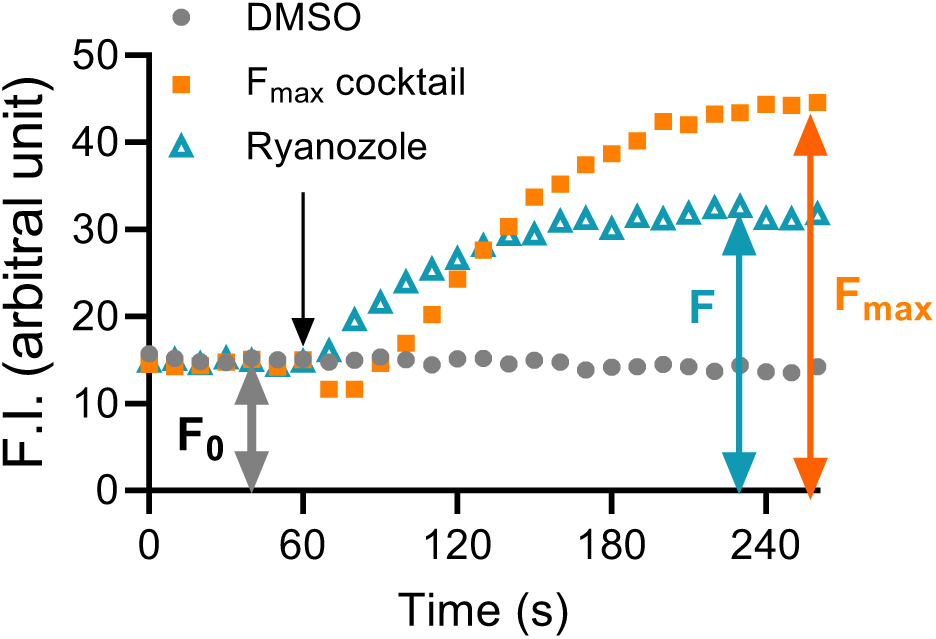
Fluorescence parameters for ER Ca²⁺ measurements. Representative time-lapse R-CEPIA1er fluorescence traces measured using FlexStation 3 fluorometer. Solutions containing DMSO (final 0.2%), test compound (1 μM Ryanozole), or an Fmax cocktail (final 20 μM ionomycin, 20 mM CaCl₂, and 10 mM caffeine) were added at 60 s (arrow). F0 was defined as the average fluorescence during the initial 60 s, whereas F and Fmax were defined as the average fluorescence during the final 60 s of the recording after addition of the test compound and the Fmax cocktail, respectively.

**Extended Data Fig. 2.**
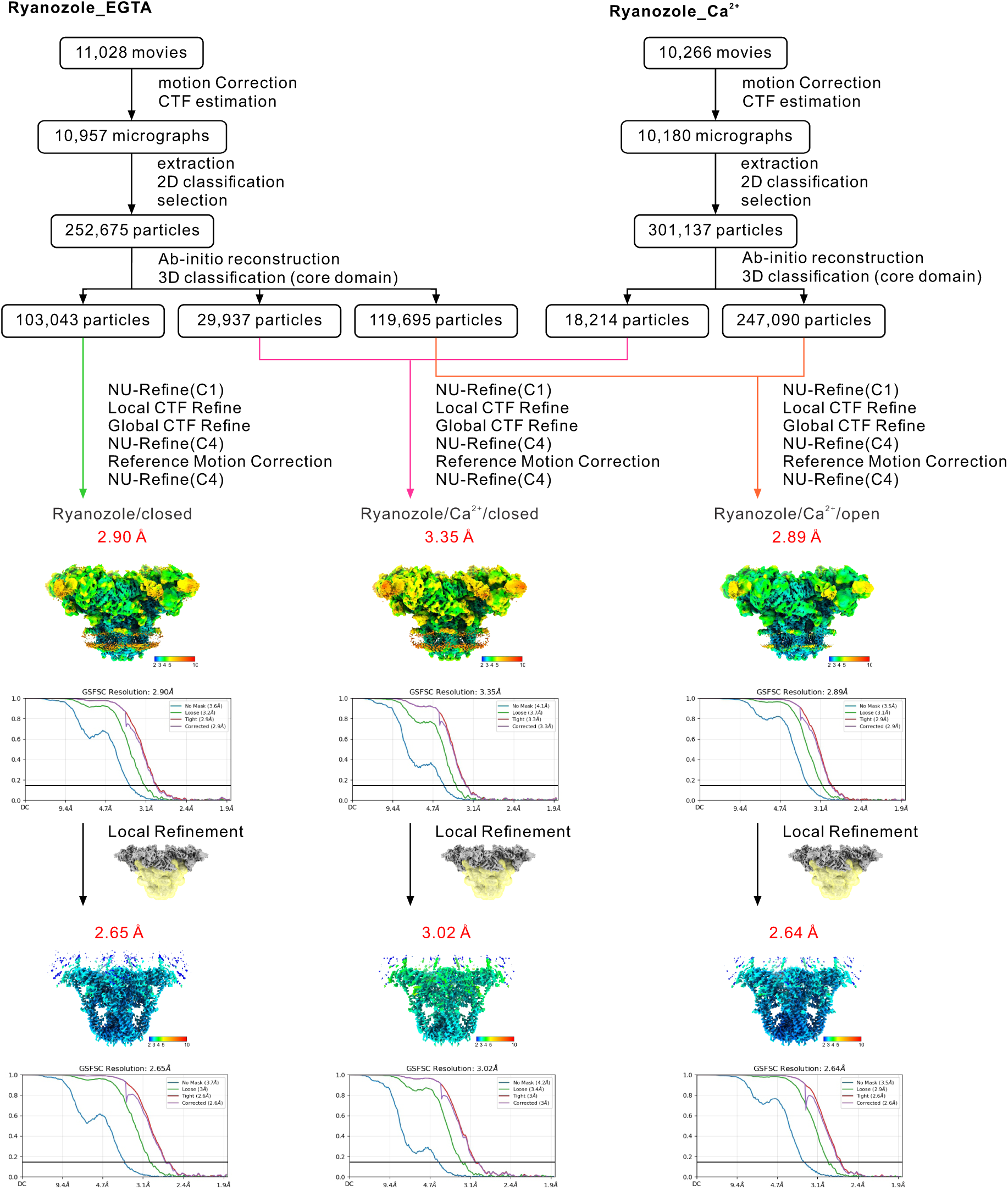
Workflows for cryo-EM data processing and estimations of resolution by Fourier shell correlation (FSC) plots and local resolution EM maps. (Left) RyR2 in the presence of EGTA and Ryanozole, (Right) RyR2 in the presence of Ca^2+^ and Ryanozole.

**Extended Data Fig. 3.**
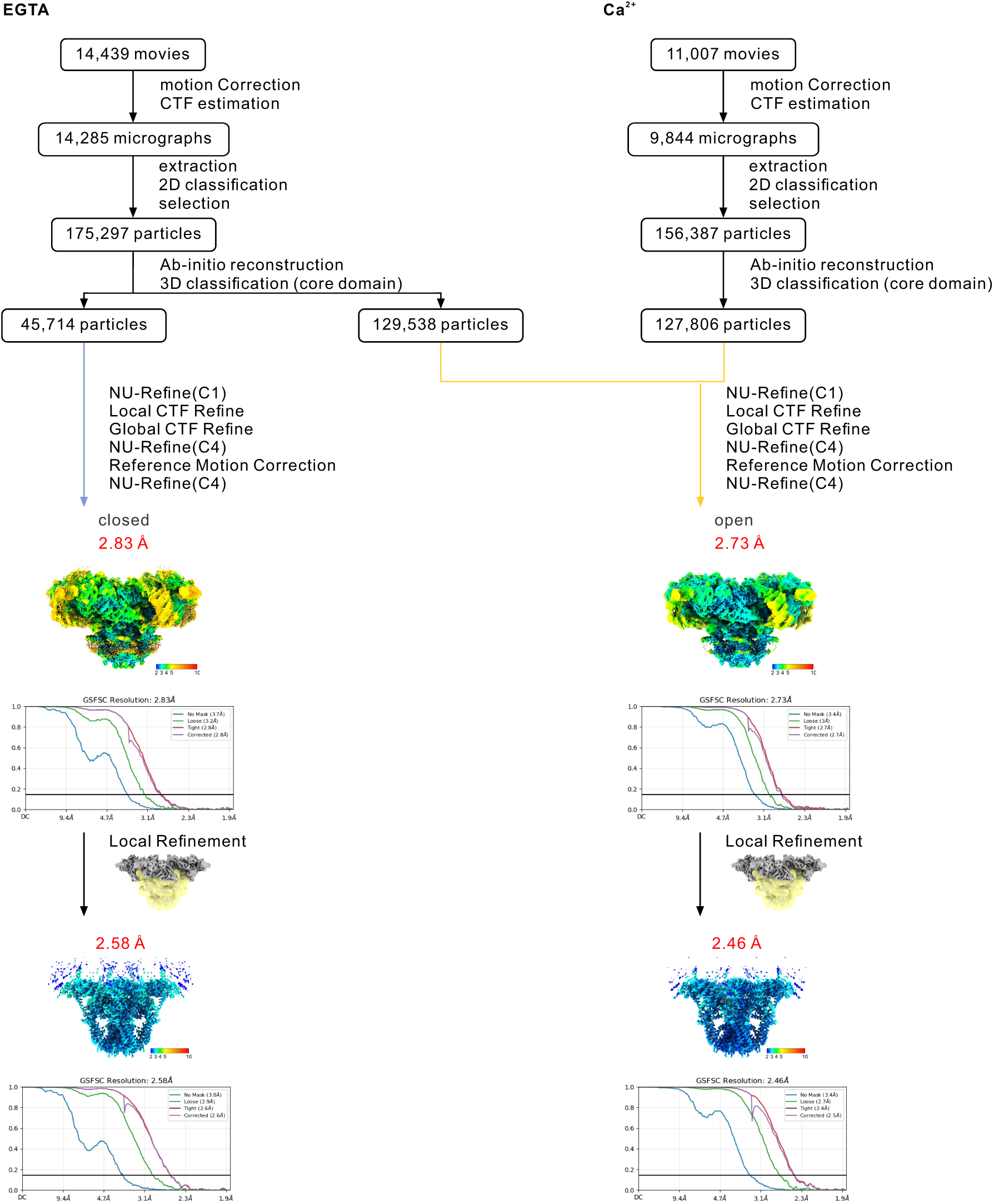
Workflows for cryo-EM data processing and estimations of resolution by Fourier shell correlation (FSC) plots and local resolution EM maps. (Left) RyR2 in the presence of EGTA, (Right) RyR2 in the presence of Ca^2+^.

**Extended Data Fig. 4.**
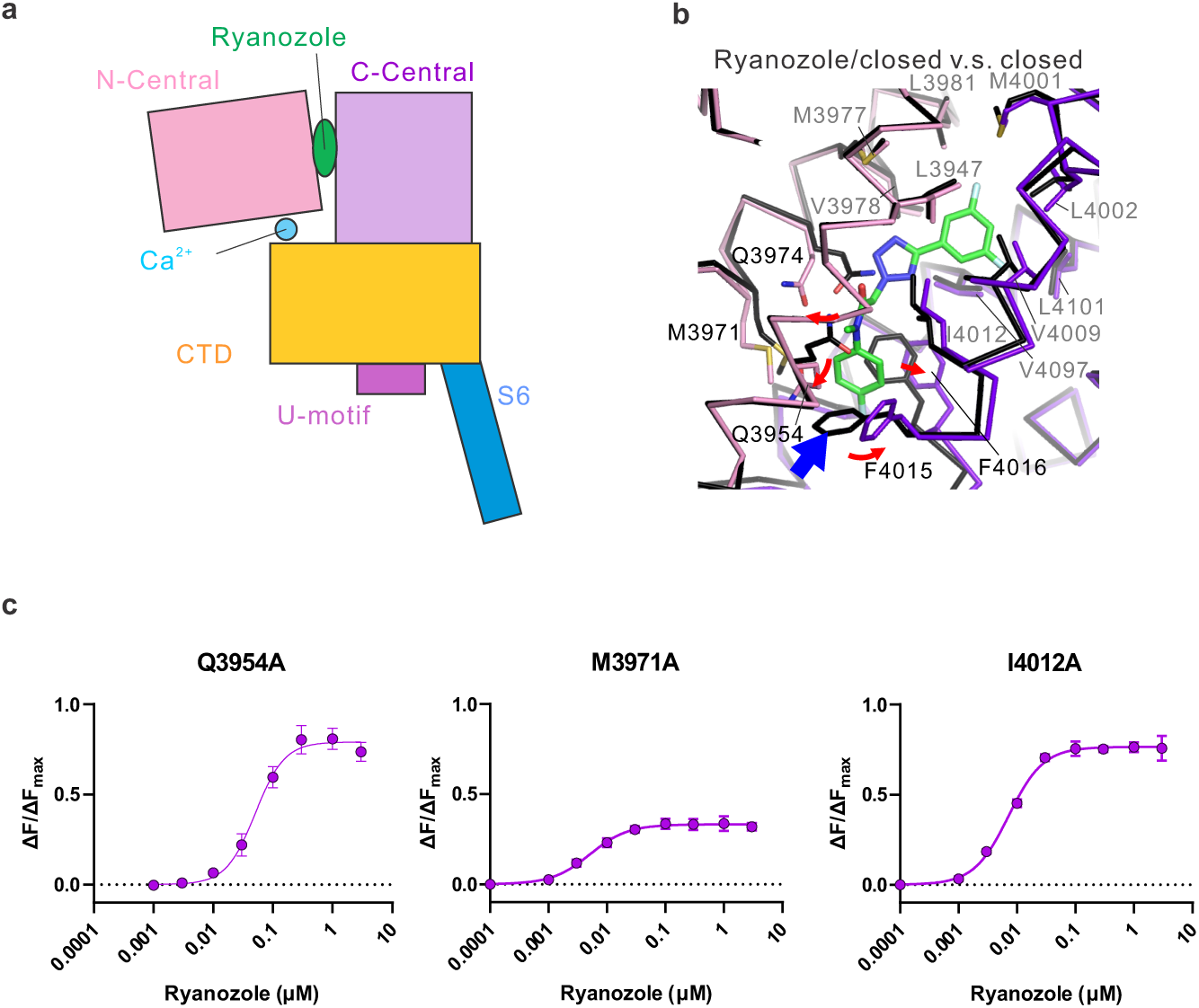
Details of Ryanozole-binding site. (a) Domain architecture of the RyR2 core domain. (b) Local structural changes upon Ryanozole binding, shown by comparison between the closed state and Ryanozole/closed conformation. Ryanozole/closed conformation is overlaid with the main chain of the closed state (black). (c) Effects of alanine substitutions near the Ryanozole-binding site. Residues showing marked alterations in the concentration-response to Ryanozole, except those shown in Fig 2d, are indicated.

**Extended Data Fig. 5.**
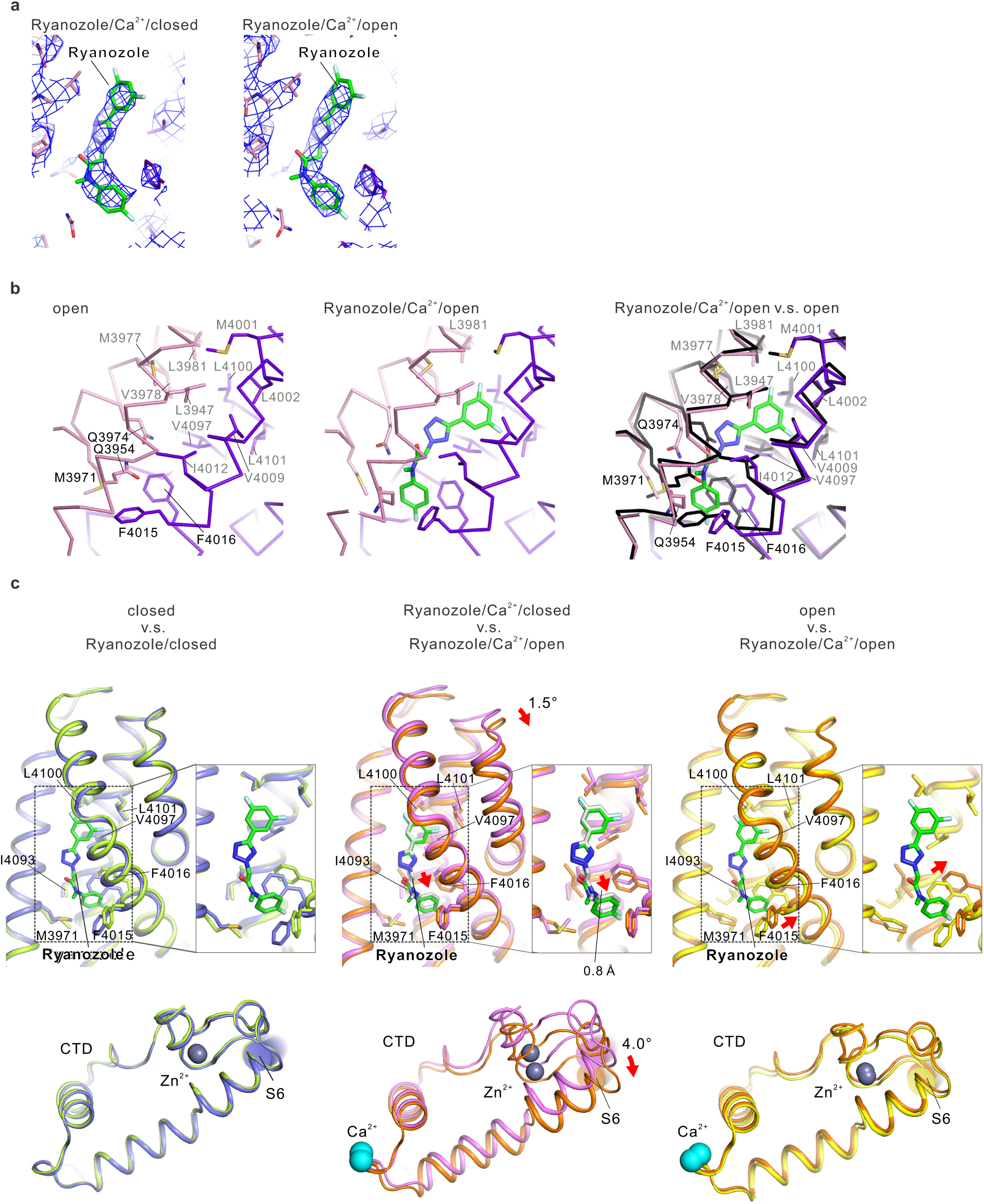
Comparison with other conformations. (a) Cryo-EM density maps of bound Ryanozole. Left, Ryanozole/Ca^2+^/closed conformation. Right, Ryanozole/Ca^2+^/open conformation. (b) Structural details near the bound Ryanozole: Left, Ryanozole-unbound open state. Middle, Ryanozole/ Ca^2+^/open conformation. Right, overlay of the Ryanozole/Ca^2+^/open with the main chain of the open state (black). The domains are colored as shown in Fig. 1e. Ryanozole is a stick model containing green carbon atoms. (c) Structural comparisons highlighting ryanzole-induced domain movement. Top: Relative movement of the C-central domain with respect to the N-central domain after the superposition of the N-central domains. The inset on the right shows a magnified view of the dashed-boxed area on the left. Bottom: CTD movement after N-central domain superposition. Left: Comparison between the closed state and the ryanozole/closed conformation. Middle: Comparison between the Ryanozole/Ca²⁺/closed and Ryanozole/Ca²⁺/open conformations. Right: Comparison between the open state and the Ryanozole/Ca²⁺/open conformation. Cα ribbons are colored for the Ryanozole-bound states and shown in black for the respective reference states.

**Extended Data Fig. 6.**
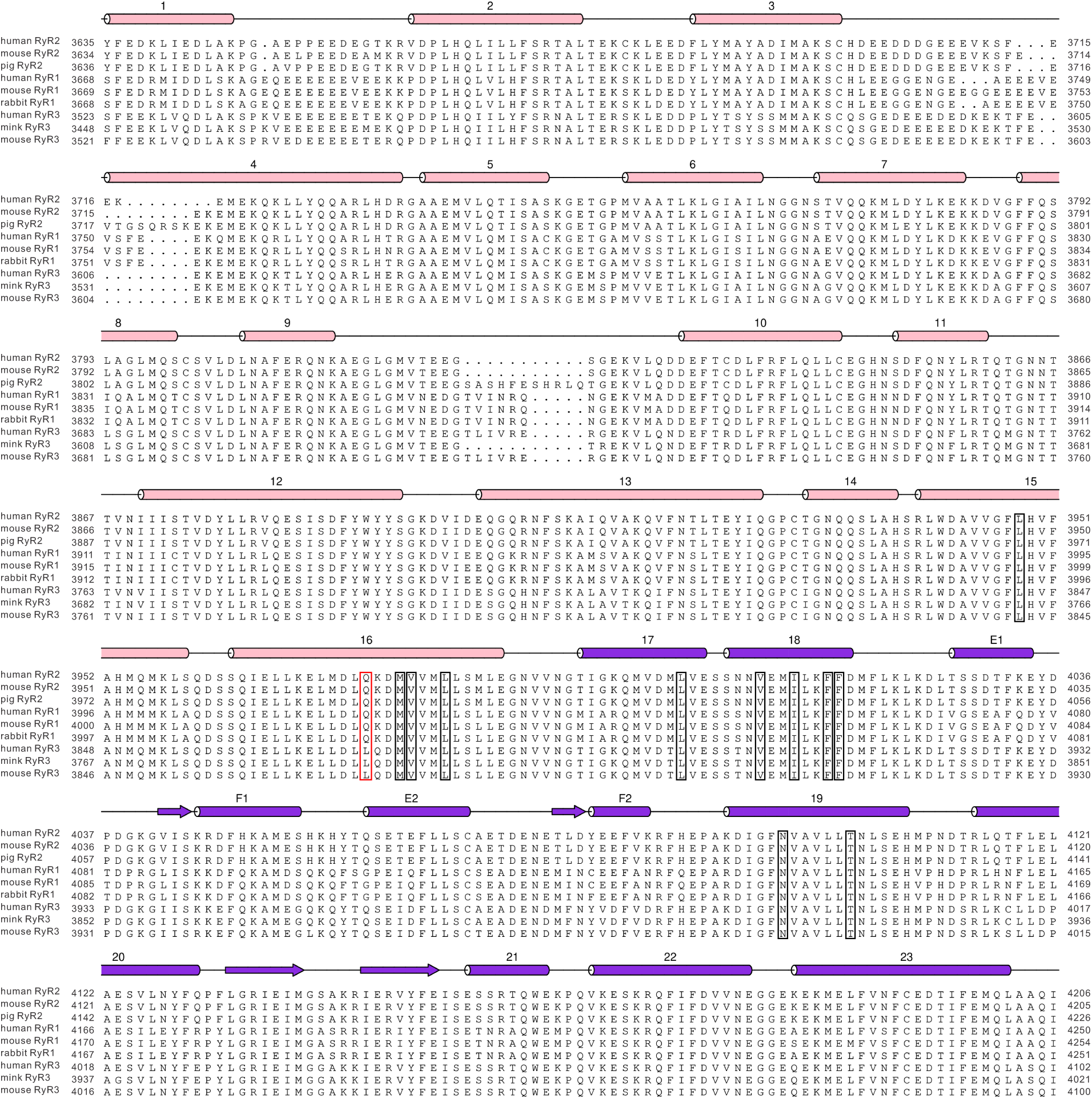
Amino acid sequence alignment of RyR isoforms near the Ryanozole-binding site in RyR2. The amino acid residues conserved across the three subtypes are indicated by black squares. Non-conserved residues of particularly functional importance are highlighted with red squares. The numbers above each square denote the corresponding amino acid positions in mouse RyR2.

**Supplementary Table 1.**
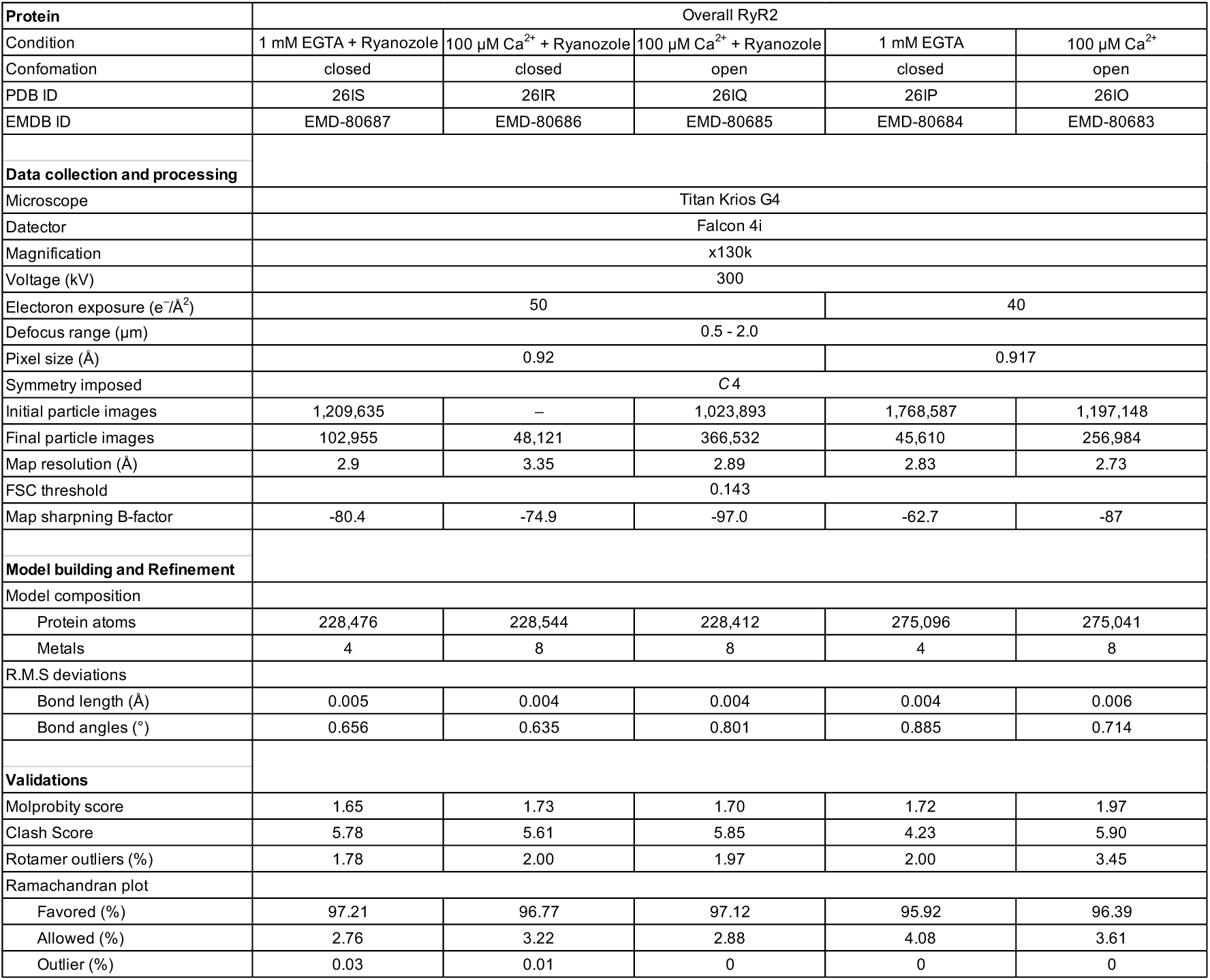
Cryo-EM data collection, refinement, and validation statistics for the overall RyR2 structure.

| Protein | Overall RyR2 |  |  |  |  |
| --- | --- | --- | --- | --- | --- |
| Condition | 1 mM EGTA + Ryanazole | 100 μM Ca<sup>2+</sup> + Ryanazole | 100 μM Ca<sup>2+</sup> + Ryanazole | 1 mM EGTA | 100 μM Ca<sup>2+</sup> |
| Conformation | closed | closed | open | closed | open |
| PDB ID | 26IS | 26IR | 26IQ | 26IP | 26IO |
| EMDB ID | EMD-80687 | EMD-80686 | EMD-80685 | EMD-80684 | EMD-80683 |
| Data collection and processing |  |  |  |  |  |
| Microscope | Titan Krios G4 |  |  |  |  |
| Detector | Falcon 4i |  |  |  |  |
| Magnification | x130k |  |  |  |  |
| Voltage (kV) | 300 |  |  |  |  |
| Electron exposure (e<sup>-</sup>/Å<sup>2</sup>) | 50 |  |  | 40 |  |
| Defocus range (μm) | 0.5 - 2.0 |  |  |  |  |
| Pixel size (Å) | 0.92 |  |  | 0.917 |  |
| Symmetry imposed | C4 |  |  |  |  |
| Initial particle images | 1,209,635 | – | 1,023,893 | 1,768,587 | 1,197,148 |
| Final particle images | 102,955 | 48,121 | 366,532 | 45,610 | 256,984 |
| Map resolution (Å) | 2.9 | 3.35 | 2.89 | 2.83 | 2.73 |
| FSC threshold | 0.143 |  |  |  |  |
| Map sharpening B-factor | -80.4 | -74.9 | -97.0 | -62.7 | -87 |
| Model building and Refinement |  |  |  |  |  |
| Model composition |  |  |  |  |  |
| Protein atoms | 228,476 | 228,544 | 228,412 | 275,096 | 275,041 |
| Metals | 4 | 8 | 8 | 4 | 8 |
| R.M.S deviations |  |  |  |  |  |
| Bond length (Å) | 0.005 | 0.004 | 0.004 | 0.004 | 0.006 |
| Bond angles (°) | 0.656 | 0.635 | 0.801 | 0.885 | 0.714 |
| Validations |  |  |  |  |  |
| Molprobrity score | 1.65 | 1.73 | 1.70 | 1.72 | 1.97 |
| Clash Score | 5.78 | 5.61 | 5.85 | 4.23 | 5.90 |
| Rotamer outliers (%) | 1.78 | 2.00 | 1.97 | 2.00 | 3.45 |
| Ramachandran plot |  |  |  |  |  |
| Favored (%) | 97.21 | 96.77 | 97.12 | 95.92 | 96.39 |
| Allowed (%) | 2.76 | 3.22 | 2.88 | 4.08 | 3.61 |
| Outlier (%) | 0.03 | 0.01 | 0 | 0 | 0 |

**Supplementary Table 2.** Cryo-EM data collection, refinement, and validation statistics for the RyR2 core domain.

| Protein | RyR2-Core domain |  |  |  |  |
| --- | --- | --- | --- | --- | --- |
| Condition | 1 mM EGTA + Ryanazole | 100 μM Ca<sup>2+</sup> + Ryanazole | 100 μM Ca<sup>2+</sup> + Ryanazole | 1 mM EGTA | 100 μM Ca<sup>2+</sup> |
| Conformation | closed | closed | open | closed | open |
| PDB ID | 26IX | 26IW | 26IV | 26IU | 26IT |
| EMDB ID | EMD-80692 | EMD-80691 | EMD-80690 | EMD-80689 | EMD-80688 |
| Data collection and processing |  |  |  |  |  |
| Microscope | Titan Krios G4 |  |  |  |  |
| Detector | Falcon 4i |  |  |  |  |
| Magnification | x130k |  |  |  |  |
| Voltage (kV) | 300 |  |  |  |  |
| Electron exposure (e<sup>-</sup>/Å<sup>2</sup>) | 50 |  |  | 40 |  |
| Defocus range (μm) | 0.5 - 2.0 |  |  |  |  |
| Pixel size (Å) | 0.92 |  |  | 0.917 |  |
| Symmetry imposed | C4 |  |  |  |  |
| Initial particle images | 1,209,635 | – | 1,023,893 | 1,768,587 | 1,197,148 |
| Final particle images | 102,955 | 48,121 | 366,532 | 45,610 | 256,984 |
| Map resolution (Å) | 2.65 | 3.02 | 2.64 | 2.58 | 2.46 |
| FSC threshold | 0.143 |  |  |  |  |
| Map sharpening B-factor | -75.8 | -74.0 | -85.8 | -60.1 | -74.5 |
| Model building and Refinement |  |  |  |  |  |
| Model composition |  |  |  |  |  |
| Protein atoms | 67,428 | 67,452 | 67,360 | 67,268 | 67,212 |
| Metals | 4 | 8 | 8 | 4 | 8 |
| R.M.S deviations |  |  |  |  |  |
| Bond length (Å) | 0.004 | 0.004 | 0.003 | 0.003 | 0.004 |
| Bond angles (°) | 0.549 | 0.558 | 0.532 | 0.552 | 0.554 |
| Validations |  |  |  |  |  |
| Molprobrity score | 1.30 | 1.32 | 1.28 | 1.42 | 1.40 |
| Clash Score | 3.83 | 3.44 | 3.77 | 3.91 | 3.50 |
| Rotamer outliers (%) | 0.86 | 1.16 | 1.08 | 1.24 | 1.67 |
| Ramachandran plot |  |  |  |  |  |
| Favored (%) | 97.34 | 97.24 | 97.58 | 97.03 | 97.58 |
| Allowed (%) | 2.66 | 2.76 | 2.42 | 2.97 | 2.42 |
| Outlier (%) | 0 | 0 | 0 | 0 | 0 |

**Supplementary Table 3.**
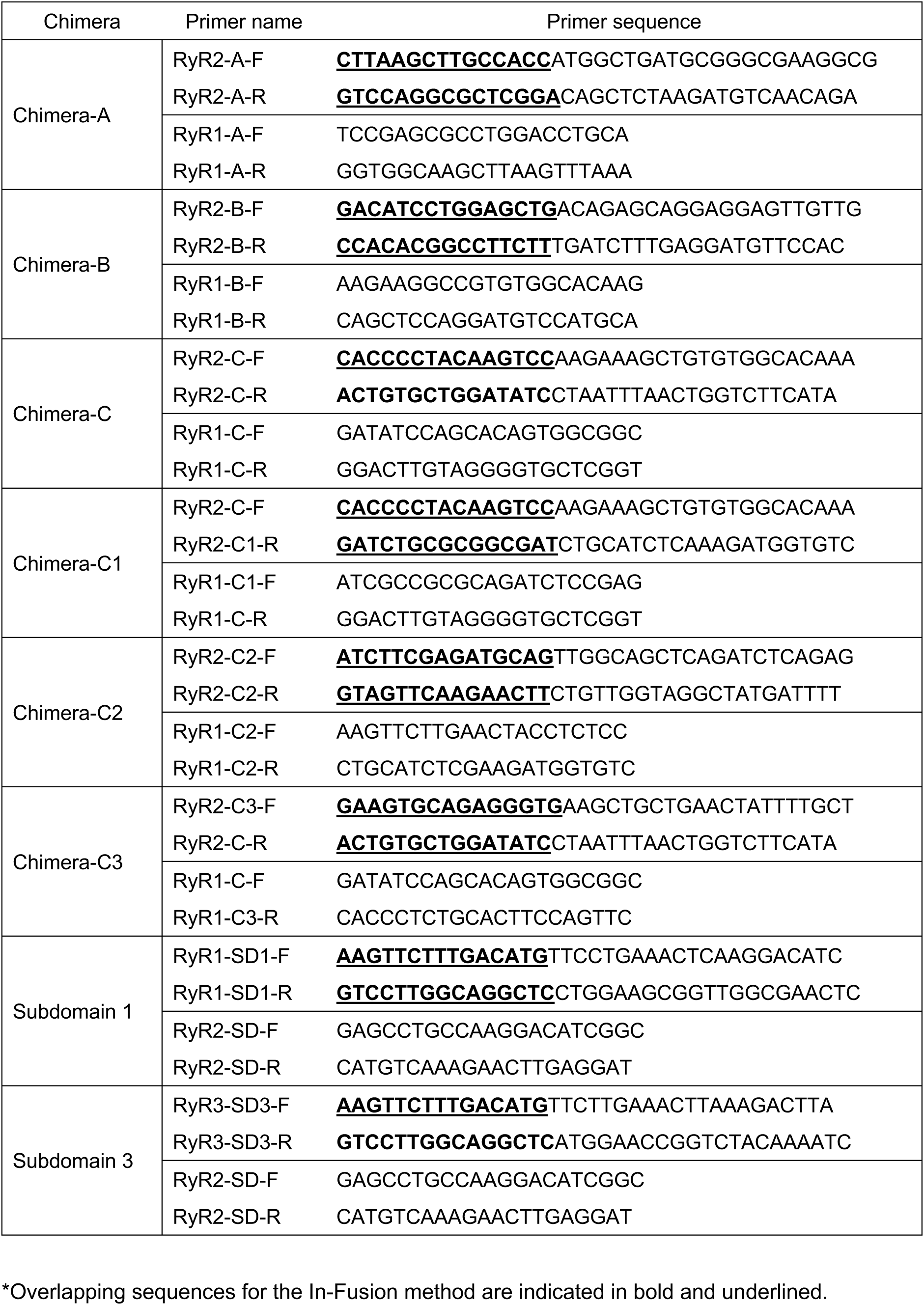
Primer sequences for the RyR1/RyR2 or RyR2/RyR3 chimera.

**Supplementary Table 4.**
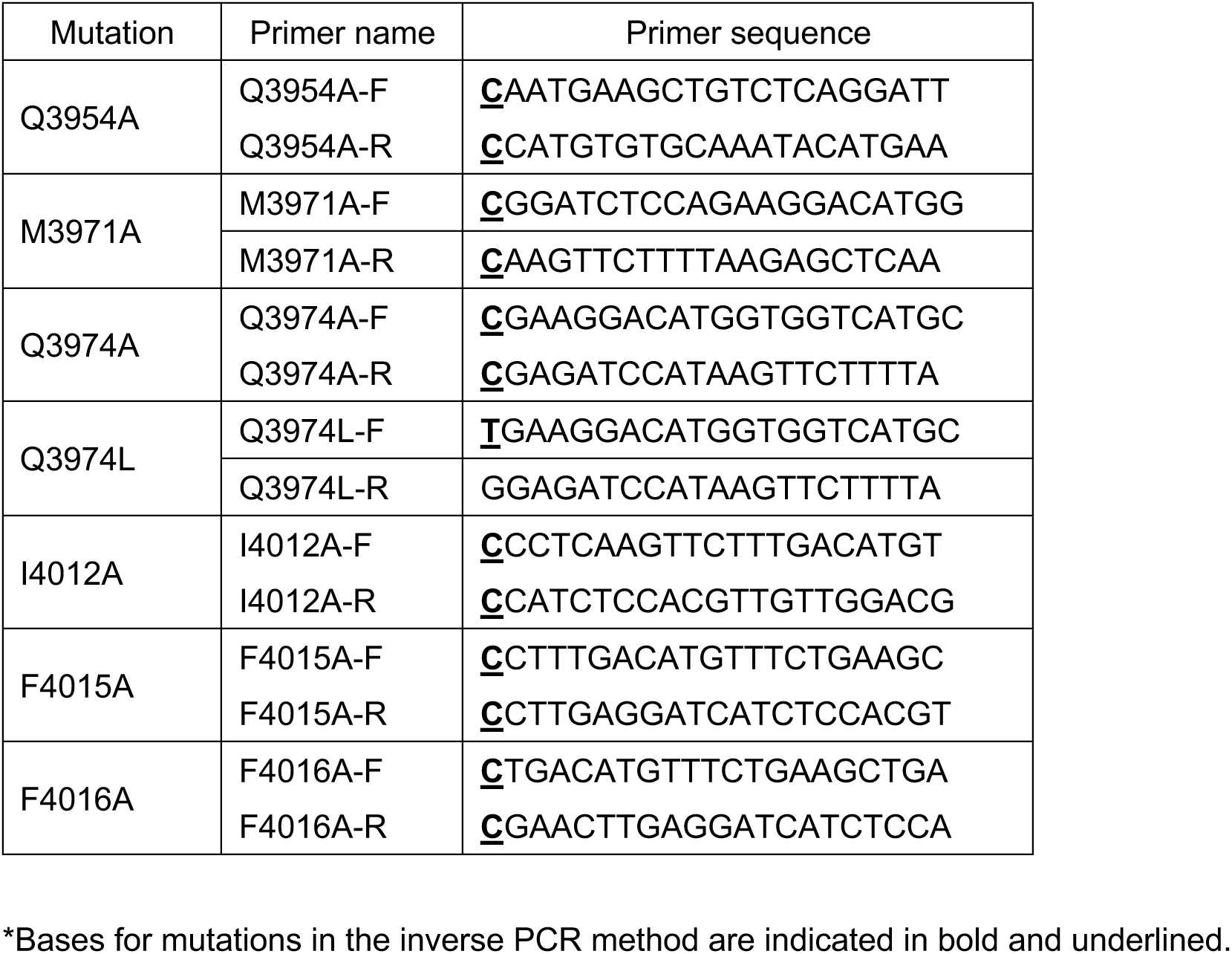
Primer sequences for the point mutations of RyR2.

**Supplementary Movie 1.** Conformational change upon Ryanazole binding (closed → Ryanazole/Ca²⁺/closed).

**Supplementary Movie 2.** Conformational shift of the C-Central domain upon Ryanozole binding.

**Supplementary Movie 3.** Conformational shift of the CTD upon Ryanozole binding.

